# Machine Learning–Guided Structure–Activity Discovery of Polymer Configurations in Lipid Nanoparticles for Kiss-and-Run Endosomal Escape

**DOI:** 10.1101/2025.08.13.670120

**Authors:** Jiamin Wu, Xia Qin, Po-Yu Chou, Binh Tran, Isaiah O. Betinol, Alex Birkenshaw, Haider Hussain, Ethan Curtis, Natalie Jones, Jolene Reid, Colin Ross, Shyh-Dar Li

## Abstract

Endosomal escape remains a major barrier to effective nucleic acid delivery via lipid nanoparticles (LNPs). Here, we address this challenge by incorporating a pH-sensitive polymer, polyhistidine, into LNPs (pLNPs) to facilitate endosomal escape, with a focus on optimizing the polymer’s molecular weight (MW) and configuration—parameters that remain largely unexplored. Through systematic engineering, we designed linear and branched polyhistidine architectures with varied MWs and configurations. *In vivo* screening identified an optimized pLNP formulation incorporating a symmetrical bis-lysine histidine dendron with a MW of ∼1800 g/mol, which achieved a 266-fold increase in liver bioluminescence following intravenous delivery of luciferase mRNA compared to standard LNPs at an equivalent RNA dose. Mechanistic studies revealed that polymer configuration within pLNPs is critical for eliciting the proton sponge effect, leading to osmotic swelling and endosomal rupture. This configuration also promoted rapid endosomal membrane destabilization via a kiss-and-run mechanism, enabling efficient cytosolic release. When delivering base editor mRNA and single-guide RNA, the optimized pLNPs achieved 8% gene editing efficiency in the mouse liver at a low dose of 0.1 mg/kg, compared to 1% with standard LNPs. To accelerate discovery and address macromolecular design challenges, we developed a machine learning (ML) framework based on amino acid-level graph neural networks (GNNs). This approach identified branched, dendritic configurations with densely arranged histidine residues on a multivalent core as key determinants of delivery performance. The top ML-predicted candidate, NS535, achieved a 705-fold increase in liver bioluminescence over standard LNPs, validating our data-driven design strategy. Together, these findings establish a closed-loop platform integrating rational design, mechanistic validation, and ML-guided optimization to advance RNA delivery. By elucidating structure-activity relationships for polyhistidine carriers and demonstrating efficient, low-dose genome editing, this work provides a blueprint for next-generation nucleic acid therapeutics.

## Introduction

Approximately 7,000–10,000 diseases are caused by single-gene mutation^1^. Base editors, which precisely and permanently modify individual DNA bases, offer a promising therapeutic strategy by targeting the root cause^2, 3^. Notably, 48% of pathogenic human single-nucleotide polymorphisms involve C-to-T or G-to-A conversions, making adenine base editors (ABEs) particularly suitable for correction^4^. Systemic delivery of ABEs via lipid nanoparticles (LNPs) holds transformative potential, especially following the success of LNP-based mRNA vaccines^5^. However, broader therapeutic use is limited by inefficient endosomal escape (1–4%)^6, 7^, which restricts transfection efficiency.

The key lipid component in LNPs is a pH-responsive ionizable lipid^8^. This lipid containing a tertiary amino group, becomes protonated in the acidic environment of the endosome, promoting ion pairing with and destabilization of the anionic endosomal membrane, and facilitating the release of cargo into the cytoplasm^9, 10^. Although significant research has been devoted to optimizing ionizable lipids to enhance endosomal escape^11-14^, relying on ionizable lipids alone remains insufficient to fully overcome this barrier. We hypothesized that incorporating more tertiary amine groups with an optimized molecular configuration through a polymer form into the LNP formulation would increase the “proton sponge” effect, suppressing the acidification in the endosome. To counteract this, the cell would pump more protons into the endosomes, which then leads to an influx of chloride ions and water, eventually resulting in osmotic swelling and ultimately endosomal rupture and cargo release^15^. While the proton sponge effect is leveraged in various cationic polymers— such as polyethyleneimine (PEI)—to facilitate intracellular delivery^16^, their permanent positive charge, high molecular weight, non-degradability, and inherent dispersity raise concerns regarding safety and reproducibility^17, 18^. Furthermore, the impact of polymer molecular weight (MW) and configuration on endosomal release are not well characterized.

In contrast, polyhistidine offers a favorable profile: it is biodegradable, can be synthesized with a defined MW and configuration using a peptide synthesizer, and contains pH-responsive imidazole groups^19^. Here we investigated the addition of terminally capped polyhistidine with defined MW, molar percentage to ionizable lipid (mol%) and spatial configuration as a fifth component to the conventional four-component LNP system to enhance endosomal escape and transfection efficiency. However, the rational design of optimal polyhistidine remains challenging due to a critical knowledge gap: the impact of structural features—such as chain length, branching patterns, and spatial configuration—on delivery performance is largely unexplored. This challenge is further exacerbated by the limitations of traditional structure–activity relationship (SAR) approaches, which often fail to account for the complex interactions between peptide topology and nanoparticle behavior in macromolecular systems^20, 21^. To address this, we developed a machine learning (ML) framework to systematically explore the design space of polyhistidine and elucidate its impact on mRNA delivery. We hypothesized that the molecular configuration of polyhistidine incorporated in LNPs critically dictates the dense, spatial arrangement of imidazole groups (pKa 6–6.5), thereby enhancing proton sponge activity across all stages of endosomal maturation—from early to late endosomes and lysosomes—by leveraging polyhistidine’s buffering capacity under increasingly acidic conditions.

To test this hypothesis, we synthesized and screened a library of polyhistidine variants with different lengths and configurations, incorporating them into standard LNP formulations at varying ratios. Their mRNA delivery efficiency was evaluated *in vivo*. Advanced imaging techniques were employed to quantitatively monitor the endosomal escape dynamics of pLNPs, including lysosomal swelling and rupture, as well as kiss-and-run behavior. We further validated the therapeutic potential of optimized pLNPs in *in vivo* genome editing applications and systematically assessed their safety profile in mice. Additionally, the ML-guided designs were confirmed through *in vivo* studies.

## Methods and Materials

### Lipid Nanoparticle (LNP) and Peptide-Lipid Nanoparticle (pLNP) Formulations

Formulations were composed of various polyhistidine (1.22-25 mol%), DLin-MC3-DMA (MC3, 25-50 mol%) (Cayman Chemical), cholesterol (38.5 mol%) (Sigma-Aldrich), 1,2-distearoyl-sn-glycero-3-phosphocholine (DSPC,10 molt%) (Avanti Polar Lipids), and DMG-PEG (1.5-2 mol%) (Avanti Polar Lipids). Polyhistidines of different lengths and confugurations were synthesized by Synpeptide (Shanghai, China), with N- and C-termini capped with acetylation and amidation, respectively. These peptide and lipid components were dissolved in ethanol at 10 mg/mL, and rapidly mixed with an mRNA solution (25 μg/mL) in a pH 4 aqueous buffer using a microfluidic T-mixer according to the established protocol^22^. An MC3 lipid-to-mRNA N/P ratio of 6:1 was maintained. The resulting formulation was then dialyzed against PBS (pH 7.4) overnight using a 10 kDa molecular weight cutoff dialysis cassette (ThermoFisher) at 25°C, followed by sterile filtration through a 0.22 μm filter. To label the formulations with DiR or DiI, 1 wt% of the dye (ThermoFisher) was spiked.

### Formulation Characterization

Formulations were characterized for particle size, polydispersity index (PDI), and zeta potential (ZP) using a Zetasizer Nano ZS (Malvern Instruments). RNA concentration and encapsulation efficiency were measured using the Quanti-iT RiboGreen RNA assay (Invitrogen).

### Cell Culture

Human embryonic kidney (HEK) 293T cells were cultured in Dulbecco’s Modified Eagle Medium (DMEM) with 10% fetal bovine serum (FBS) and penicillin-streptomycin (100 units/mL penicillin, 100 μg/mL streptomycin; Gibco). Cells were incubated at 37°C with 5% CO□and sub-cultured up to 15 passages.

For particle tracking, cells were seeded in a 35-mm confocal dish (VWR) at a density of 2 × 10□cells/dish and incubated for 18–20 hours. Live cells were first stained with Hoechst 33342 for nuclei and Lysotracker for endo-lysosomes according to the manufacturer’s protocols. Next, 20 μg/mL (total lipids) DiI-labeled formulations were added to the cells, followed by immediate imaging by a spinning disk PerkinElmer confocal microscope (SDCM) over a 45-minute interval. Videos were analyzed with Trackmate, Matlab and ImageJ.

### Endosomal Buffering Capacity

The buffering capacity of formulations was assessed by the acid titration method. Formulations (0.4 mg/mL of total lipids in PBS, pH 7.4) were titrated with 0.1 M HCl in 5 μL increments, and pH changes were monitored with a pH meter.

### XTT Assay for Cell Viability

HEK293T cells were seeded at a density of 5 × 10□cells/well in a 96-well plate and incubated for 24 hours (37°C, 5% CO□, humidified). Cells were then treated with various concentrations of formulations. After 24 hours, cell viability was evaluated using an XTT assay as per standard protocols^23^.

### Cell membrane destabilization assay

Freshly drawn murine whole blood was processed to isolate erythrocytes through centrifugation (10,000g, 5 min). The obtained red blood cell (RBC) fraction underwent five successive washing cycles using phosphate-buffered saline (PBS, pH 7.4). Washed RBCs were then resuspended in PBS adjusted to either pH 7.4 or 5.5 for subsequent testing. For the compatibility assay, RBC suspensions were distributed into 96-well plates and exposed to either LNPs or pLNPs at total lipids concentration at 0.15, 0.3, 0.6 and 0.12 mM. Following a 60-minute incubation period at 37°C, samples were centrifuged (10,000g, 5 min) to separate cellular components from supernatant. Hemoglobin release was quantified spectrophotometrically by measuring absorbance at 540 nm. Control samples included: (1) baseline hemolysis (RBCs in PBS only) and (2) complete hemolysis (RBCs in 1% w/v Triton X-100 solution).

These controls established the reference ranges for minimal and maximal hemoglobin release, respectively.

### Endosome-mimicking membrane disruption assay by fluorescence resonance energy transfer (FRET) assay

The membrane disruption activity of different formulations was evaluated using a fluorescence resonance energy transfer (FRET) approach. Synthetic liposomes were formulated to simulate endosomal membrane by combining DOPS, DOPC, DOPE, NBD-PE (Avanti Polar Lipids), and Rho-PE (Avanti Polar Lipids) in a 25:25:48:1:1 molar ratio. The lipid mixture in chloroform was processed through rotary evaporation followed by 2 hours of vacuum drying to create a thin lipid film. This film was then rehydrated in phosphate-buffered saline (pH 7.4) and subjected to 30 minutes of sonication, yielding a 1 mM lipid suspension. For the fusion assay, different pH conditions (5.5 or 7.4) were established in a black 96-well plate with 100 μL PBS per well. Each well received 5 μL of the prepared anionic liposome suspension along with 10 μL of either LNP or pLNP test samples. Following a 5-minute incubation at 37°C, fluorescence intensity was measured at 465 nm excitation and 520 nm emission wavelengths. Control measurements included: Baseline fluorescence (Fmin) from liposomes in PBS alone; Maximum fluorescence (Fmax) from liposomes treated with 2% Triton X-100. Membrane disruption efficiency was quantitatively determined using the formula: [(F - Fmin)/(Fmax - Fmin)] × 100%, where F represents the sample fluorescence reading.

### Animal Studies

Ten-to-twelve-week-old male and female CD-1 mice were acquired from Charles River Laboratories. All *in vivo* experiments were conducted under the University of British Columbia’s Animal Care Committee-approved protocol (A22–0141).

### *In Vivo* Screening of pLNP Formulations Containing Different Polyhistidines for Luciferase mRNA Delivery

Five hours post-intravenous injection (i.v.) of formulations, mice were administered 0.15 mL D-luciferin potassium salt (Cayman Chemical) (20 mg/mL in PBS) intraperitoneally (i.p.). Mice were then anesthetized under 2.5% isoflurane through inhalation, and bioluminescence imaging was performed 10 minutes post-injection using an *in vivo* imaging system (IVIS, LagoX). Image acquisition and quantification were performed with the Aura Live Image software. For bioluminescence, the exposure time was 120 s (medium binning, f-stop 1), and for fluorescence imaging, the exposure time was 20 s (medium binning, f-stop 1) with an excitation wavelength of 754 nm and an emission filter at 778 nm. Sample size was 3 per group (10-12 weeks).

### Intracellular Distribution of mRNA and Particles Within the Liver

DiI-labeled particles encapsulating FITC-mRNA were injected i.v. to mice. Five hours post-injection, mice were euthanized, and the liver was collected, embedded in OCT, and sectioned at 10 μm on a Leica Cryostat. The tissue section was fixed in 10% formalin for 10 minutes, permeabilized with 0.1% Triton X-100 for 5 minutes, and blocked with 1% BSA in PBST (PBS + 0.1% Tween 20) for 60 minutes. Following washes, the section was incubated with a primary antibody (5 μg/mL APPL1 (ThermoFisher), 2 μg/mL EEA1 (ThermoFisher), 1:100 diluted Rab11 (ThermoFisher), or 1:100 diluted Anti-LBPA (Sigma-Aldrich)) for 1 hour, followed by 3 times of PBS washes. The secondary antibody (Goat anti-Rabbit/Mouse IgG Alexa 647-conjugated (Biolegend), 1 μg/mL) was applied on the section for 1 hour at room temperature, which was washed for 3 times and stained with DAPI (Sigma-Aldrich) and were then imaged by confocal laser scanning microscope (CLSM) and analyzed with ZEN software (Carl Zeiss, Oberkochen, Germany) and image J.

### *In vivo* base editing analysis

Base editor mRNA was obtained from TriLink, while single guide RNA (sgRNA) was sourced from either Synthego or Integrated DNA Technologies (IDT). Mice received i.v. of LNP-formulated RNA at a total RNA dose of 0.1 mg/kg (mRNA: sgRNA=1:1, w/w). Both male and female mice were included in the study (2 females and 1 male per group). D-luciferin was freshly prepared in PBS at a concentration of 15 mg/mL and sterilized using a 0.2 μm filter. For *in vivo* imaging, mice were administered D-luciferin i.p. at 150 mg/kg, 10 minutes prior to imaging with the IVIS Lumina II system using a 120-second exposure time. Following imaging, mice were euthanized via deep isoflurane anesthesia followed by cervical dislocation. Organs were collected and rapidly frozen in liquid nitrogen. Tissue samples were subsequently homogenized using a FastPrep homogenizer (MP Biomedicals) at speed setting 6 for 20 seconds, repeated three times. The homogenates were then processed for Sanger sequencing analysis.

### *In Vivo* Safety Test

Mice received the formulation at 0.3 mg/kg i.v.. After 3 hours or 24 hours, animals were euthanized, and blood and serum samples were collected for analysis of inflammatory cytokines including IL-6 and TNF-α by ELISA (ThermoFisher) or safety biomarkers, including but not limited to urea (BUN), ALT, AST, ALP, creatine kinase and osmolality.

### Machine learning model

We compiled experimental data from pLNPs, incorporating polyhistidine structures, peptide mol%, and bioluminescence intensities at various time points. Each histidine peptide was featured as described above and fed into a graph neural network (GNN) with three graph convolution layers based on the GNN operator from Morris et al^24^ (Formula 1).

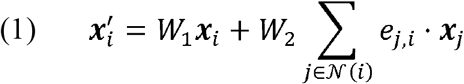

Where *x*_*i*_ is a feature vector of node *i, x*_*j*_ is a feature vector of neighbouring node *j*, ***e***_*j,i*_ is the edge weight from node *j* to node *i*, *N*(*i)* is the set of neighbours of node *i, w*_l_ and *w*_2_ are learnable weight matrices, and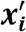 is the updated feature vector of node *i*. After each convolutional layer, a ReLU activation function is applied. Following the three graph convolution layers, node embeddings are aggregated using global mean pooling, which averages node features for final graph-level predictions (Formula 2).

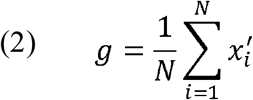

Where *N* is the total number of nodes in the graph and *g* is the graph-level representation. As the maximum radiance was dependent on external factors (peptide loading and time), these additional graph-level features were concatenated with the graph representation before being passed through a multi-layer perceptron (MLP) with one hidden layer to provide the final prediction.

### Statistical Analysis

Data were analyzed using GraphPad Prism 9 (GraphPad Software). Results are presented as mean ± s.d. * denotes *P*□<□0.05, ** denotes *P*□<□0.01, *** denotes *P*□<□0.001 and **** denotes *P*□<□0.0001.

## Results and discussion

### *In Vivo* Screening to Determine the Optimal Length of Polyhistidine for pLNP Formulation

The polyhistidine and formulation coding logics are indicated in Suppl. Figure 1 and Suppl Table 1. Linear polyhistidines of different lengths (acetyl-H_4_-amide to acetyl-H_40_-amide, coded as NS304 to NS340) (Figure 1A) were incorporated into standard MC3-based LNP at a range of mol% (2.38-25 mol%) to prepare different peptide-LNPs (pLNPs). As shown in Suppl. Table 2, vast majority of pLNPs containing acetyl-H_4_-amide to acetyl-H_14_-amide (NS304 to NS314) maintained an average size of ∼85 nm with a polydispersity index (PDI) <0.3 that were comparable to the standard LNP (MC-000-101). Incorporating longer polyhistidines (NS315 to NS340) tended to form larger particles (up to 6 μm), likely due to aggregation caused by increased interaction between longer polyhistidines and mRNA. Increasing the polyhistidine molar ratio tended to increase the particle size and decrease the mRNA encapsulation efficiency (EE%). For example, the particle size increased from 100 nm to 245 nm, and the EE% decreased from 83% to 68% when the mol% of NS313 increased from 2.38 mol% to 25 mol%. Too much polyhistidine could affect the incorporation into the LNP formulation, leading to increased particle size and decreased mRNA encapsulation. Notably, only pLNP formulations containing ≤8.33 mol% polyhistidine displayed EE% greater than 60%, among them, NS309 to NS313 displayed EE% greater than 70%, while pLNP formulations prepared outside these conditions generally exhibited EE% below 60%. Furthermore, pLNP formulations prepared with >8.33 mol% of larger molecular weight polyhistidines (NS316 to NS340) displayed aggregations and were excluded from *in vivo* screening.

**Figure 1.**
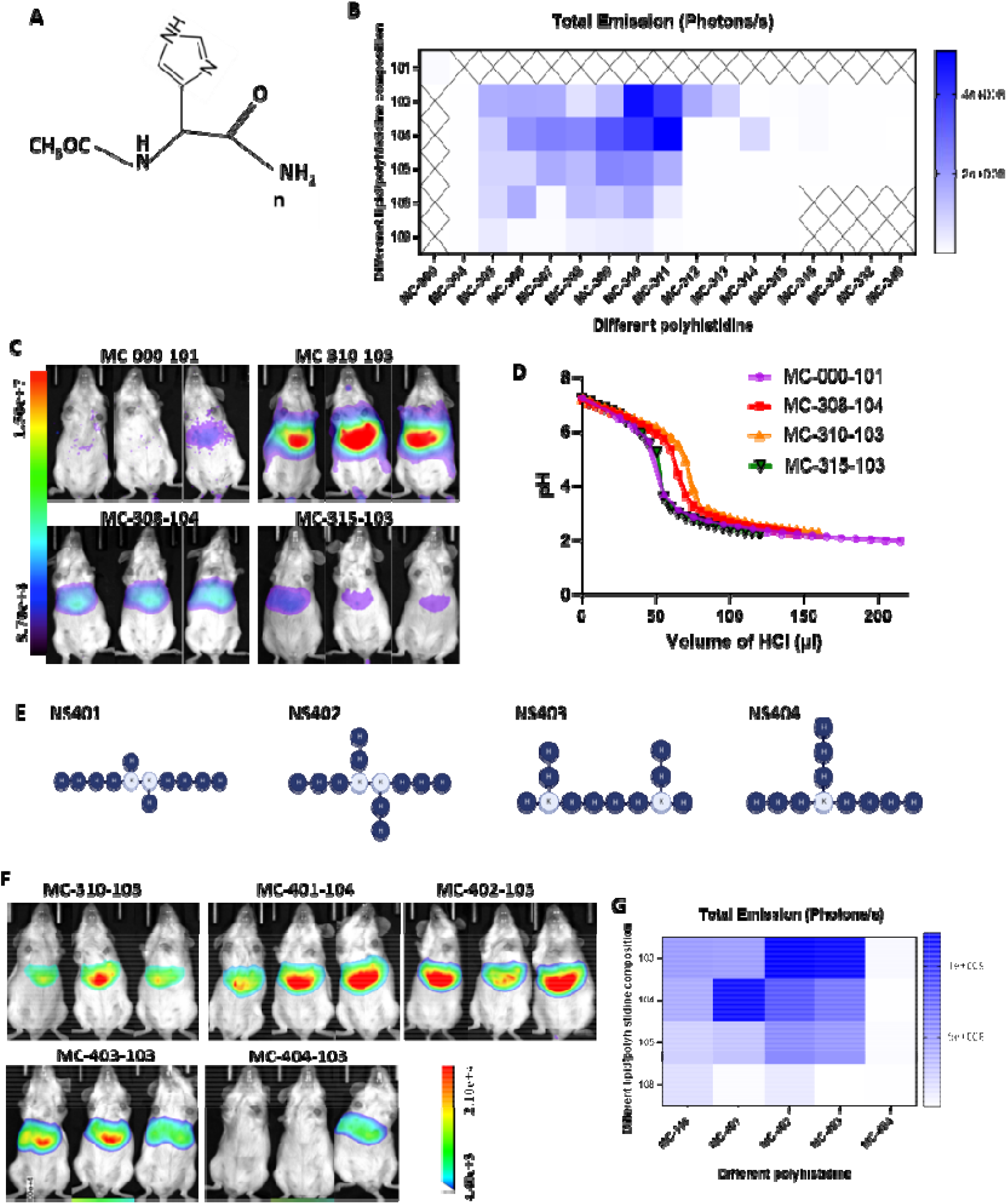
Developing and screening polyhistidine-lipid nanoparticle (pLNP) formulations. **A**. Structure of linear polyhistidine with capping at both N and C termini. **B**. Total flux of bioluminescence in the liver 5 h after i.v. delivery of different LNP and pLNP formulations loaded with luciferase mRNA (n=3). X-axis represents different polyhistidines (refer to Suppl. Figure. 1 for their structures) used to prepare pLNP formulations. Y-axis represents different lipid/polyhistidine compositions (refer to Suppl. Table 1 for compositions). **C**. Represented images of whole-body bioluminescence images of mice 5 h after i.v. delivery of different pLNPs. **D**. pH buffering effect of different linear polyhistidine incorporated pLNP formulations measured by the acid titration assay. **E**. Structures of branched polyhistine(10) (i.e., H_10_), H and K represent histidine and lysine, respectively. **F**. Represented images of whole-body bioluminescence images of mice 5 h after i.v. delivery of different pLNPs. **G**. Total flux of bioluminescence in the liver 5 h after i.v. delivery of different pLNP formulations loaded with luciferase mRNA (n=3). X-axis represents different polyhistidines (refer to Suppl. Figure. 1 for their structures) used to prepare pLNP formulations. Y-axis represents different lipid/polyhistidine compositions (refer to Suppl. Table 1 for compositions).

Since *in vitro* data do not always correspond to *in vivo* results, we screened these pLNP formulations directly in mice (n=3). Luciferase mRNA was encapsulated into different LNP and pLNP formulations and injected i.v. into mice. Mice were imaged for luciferase bioluminescence 5 hours post injection. As shown in Figure 1B and C, only formulations incorporated with NS305 to NS311 exhibited enhanced gene transfection efficiency compared to MC-000-103, and the mRNA delivery was focused to the liver. Among these, pLNPs containing NS309 to NS311 appeared optimal, showing >50-fold increased luciferase bioluminescence in the liver compared to MC-000-101. In particular, the pLNP containing 2.38 mol% NS310 (MC-310-103)was among the most potent formulations, increasing the luciferase bioluminescence in the liver by ∼90-fold compared to MC-000-101. Interestingly, the transfection efficiency decreased as the mol% of NS310 increased, likely due to the decreased EE%. These results confirmed that the MW and mol% of polyhistidine—when used as the fifth component in LNPs—are critical determinants for boosting mRNA delivery efficiency *in vivo*.

We randomly selected four LNP and pLNP formulations that displayed a wide range of transfection efficiency (Figure 1C), and compared their pH buffering effect (Figure 1D). There appeared to be a close association between the transfection efficiency and pH buffering effect: formulations with increased pH buffering effect displayed enhanced mRNA delivery efficiency, suggesting that the pH buffering effect contributed by the peptide component was a key driver of the improved transfection.

It is noted that both the N and C termini in the polyhistidines were blocked to ensure that the peptide would not become ionized after pH neutralization to 7.4. Ionized peptide could decrease the encapsulation of mRNA and increase the toxicity of the formulation due to the negative charge of the carboxylate groups that decrease the binding with mRNA and the cationic amino groups that could trigger non-specific binding with serum proteins and cell membranes, respectively. Indeed, pLNP prepared with unblocked H_10_ peptide (NS-310n) was inferior to that prepared with capped H_10_ peptide (NS-310) in mRNA encapsulation efficiency (83.9% vs 37.6%, Suppl. Table 3) and the transfection efficiency in mice (decreased by 5.5-fold, Suppl. Figure 2).

### pLNPs incorporated with branched H_10_ displayed increased *in vivo* mRNA transfection compared to linear H_10_

We then hypothesized that the configuration of polyhistidine (linear vs branched) would affect the transfection efficiency of the resulting pLNPs. We designed four branched H_10_ structures NS401–NS404, to test this hypothesis (Figure 1E). DLS analysis confirmed that all 16 pLNP formulations prepared with these branched H_10_ displayed comparable size (70-100 nm), PDI (<0.25), and zeta potential (close to neutral). When pLNPs containing < 8.33 mol% of NS401 to NS403 were used, they displayed mRNA EE >70% (Suppl. Table 4). Formulations containing NS404 had relatively low mRNA EE% (61% in MC-404-103 and 51% in MC-404-105). *In vivo* screening revealed that the transfection efficiency of the pLNP formulations was associated with the polyhistidine configuration as well as the mRNA EE%. Three out of the four branched H_10_ enhanced mRNA transfection compared to linear H_10_, with the NS402 exhibiting the highest potency (Figure 1F, G).

In particular, MC-402-103 showed comparable particle properties as MC-310-103 in particle size (80 to 90 nm), PDI (<0.15), zeta potential (∼0 mV), and mRNA EE (> 80%). Bioluminescence quantification in the liver at 5 hours post-administration showed that MC-402-103 increased the luciferase bioluminescence by 2-fold and 266-fold compared to MC-310-103 and MC-000-101, respectively (Figure 1F, G). NS404 did not enhance the transfection efficiency, likely due to its configuration and/or the decreased EE%. Similar results were obtained with H_8_ (Suppl. Figure 3 and Suppl. Table 5), showing that pLNPs prepared with branched H_8_ showed up to 2.3-fold increased transfection efficiency compared to that spiked with linear H_8_. While MW and mol% optimization established baseline activity, the configuration of polyhistidine emerged as the most critical factor governing the delivery efficiency. The branched H_10_ and H_8_ configuration demonstrated superior performance compared to their linear counterparts respectively, despite having comparable MW and incorporation mol%. This configuration-dependent activity suggests that three-dimensional arrangement of imidazole groups, rather than simply their quantity, dictated the proton sponge efficacy and membrane interactions. Again, mRNA transfection efficiency of the formulations appeared to be related to their pH buffering effect: MC-402-103 incorporating branched NS402 that displayed increased mRNA delivery efficiency had a superior pH-buffering effect compared to its counterpart formulation MC-310-103 containing linear NS310 (Suppl. Figure 4).

We then focused on comparing the optimized pLNP formulation (MC-402-103) with the standard LNP (MC-000-101) in their mechanisms of mRNA delivery.

### *In vivo* distribution, mRNA release and transfection of LNP and pLNP

To further elucidate the mechanism of the superior gene transfection efficiency of MC-402-103 over MC-000-101, we compared their *in vivo* distribution. As shown in Figure 2A, both MC-000-101 and MC-402-103 showed significant liver and spleen accumulation with minor uptake in other tissues. While there was no significant difference in the liver uptake, the accumulation of MC-402-103 in the spleen and lungs was 2.3-fold and 1.6-fold higher, respectively, than that of MC-000-101 (Figure 2B). However, only the liver displayed increased mRNA expression with MC-402-103 treatment compared to MC-000-101 (Figure 1).

**Figure 2.**
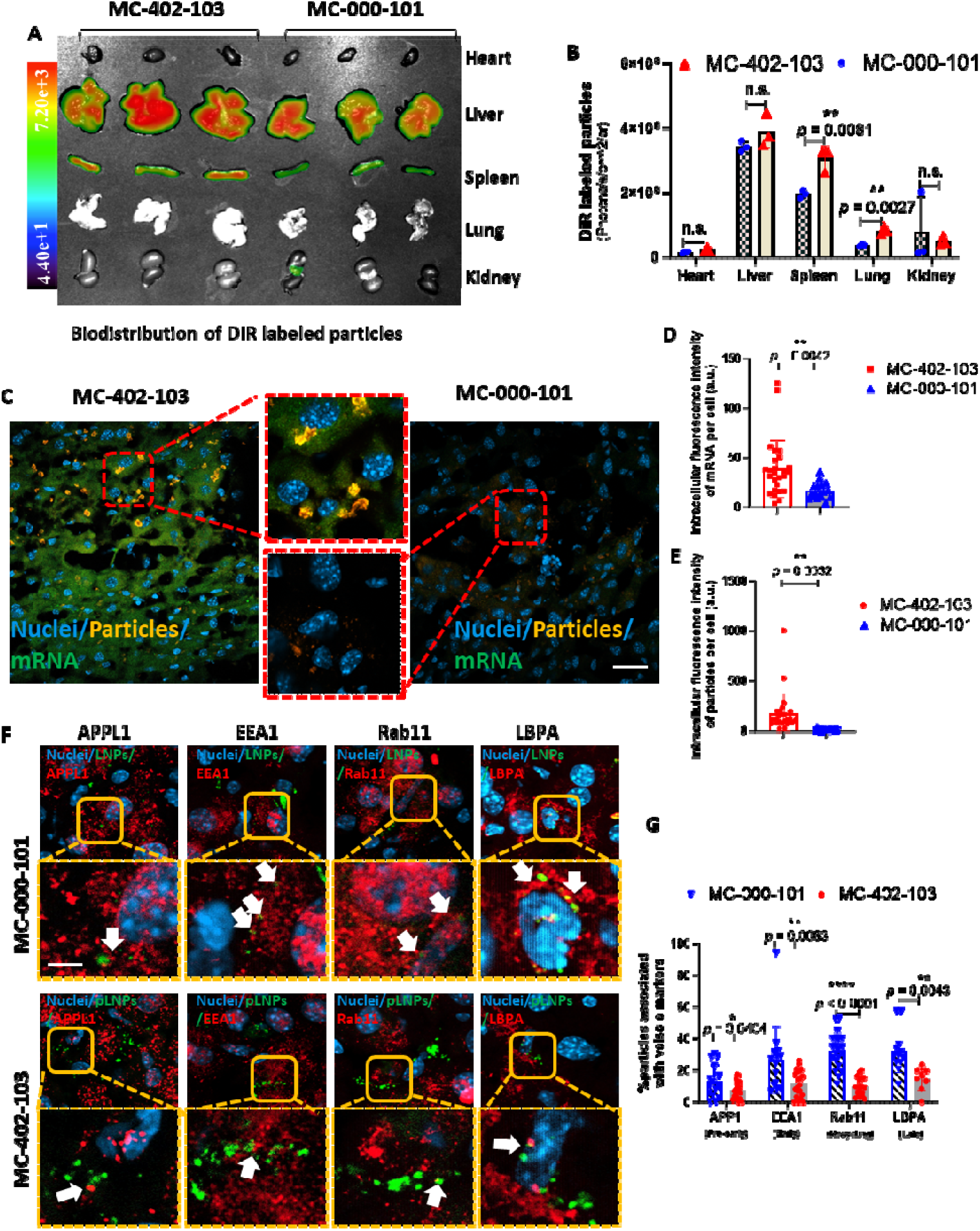
*In vivo* biodistribution and intracellular endosomal trafficking of MC-402-103 and MC-000-103 in mouse liver. Biodistribution (**A**) of DiR labeled particles and the quantitative results (**B**) in main organs in mice 5 h after i.v. injection (n=3). **C**. Intracellular uptake of DiI-label particles (yellow) loaded with FITC-mRNA (green) in the liver of mice 5 h after i.v. injection. Nuclei were stained with DAPI (blue). Scale bar = 20 µm. Quantitative results of intracellular DiI-label particles (**D**) and FITC-mRNA (**E**). **F**. Livers were collected from mice 5 h after receiving an i.v. dose of DiI-labeled particles. Liver sections were imaged using confocal microscopy. Yellow squares indicate the high-magnification view on the right. White arrows indicate association of endosomal markers with DiI particles. Nuclei were stained with DAPI (blue), MC-402-103 and MC-000-103 were labeled with DiI (green), and endosomal markers were stained with Alexa 647-antibodies (red). Scale bar = 5 µm. **G**. Quantification of % particles associated with vesicles markers. Statistical significances determined using two-tailed unpaired *t*-test (95% CI). Data are presented as mean ± SD, \**P*□<□0.05, \*\**P*□<□0.01, and \*\*\*\**P*□<□0.0001, n.s. no significant difference.

We then further analyzed the intracellular uptake of DiI-labeled MC-000-101 and MC-402-103 loaded with FITC-mRNA within the liver. Figure 2C, D revealed 500-fold higher intracellular uptake of MC-402-103 compared to MC-000-101, and MC-402-103 tended to form large clusters within the cells. Intracellular FITC-mRNA distribution analysis showed that, in the MC-402-103-treated liver, mRNA was evenly distributed within the cells, whereas barely any mRNA signal was detected in the MC-000-101-treated liver. Quantitative analysis of intracellular FITC-mRNA (Figure 2E) revealed an 8-fold increase in the MC-402-103-treated liver compared to MC-000-101. Taken together, the data in Figure 2 A to E suggested that, although effectively accumulating in the liver, MC-000-101 was largely located extracellularly, which could be washed out during the tissue section preparation. The results were consistent with the low gene transfection efficiency of MC-000-101. While the liver accumulation was comparable between MC-000-101 and MC-402-103, MC-402-103 was more effectively internalized by or retained within the liver cells, leading to increased cytosolic release of mRNA and gene expression.

### MC-402-103 exhibited reduced endosomal association among different internalization stages compared to MC-000-101 *in vivo*

We hypothesized that the symmetrical histidine dendron configuration as well as imidazole groups in NS402, with a pKa of 6-6.5, would facilitate proton absorption in acidic endosomes following the endocytosis of MC-402-103. This would enhance the pH-buffering effect (supported by Figure 1D and Suppl. Figure 4), leading to endosomal swelling and rupture through the proton sponge effect^16^. Consequently, MC-402-103 was expected to exhibit reduced association with endosomal markers and improved cytosolic release compared to MC-000-101.

To test this hypothesis, we stained four specific endosomal biomarkers and assessed their colocalization with DiI-labeled MC-000-101 or MC-402-103 following i.v. administration in mice. Livers were harvested 5 hours post-injection, cryo-sectioned, stained with Alexa 647-conjugated antibodies recognizing different endosomal markers, and analyzed via confocal microscopy. The stained markers included: APPL1, which labels very early endosomes—the first compartment post-endocytosis; EEA1, which marks early endosomes involved in sorting internalized cargo in a mildly acidic environment; Rab11, a marker of recycling endosomes that return components to the plasma membrane; and LBPA, which labels late endosomes or multivesicular bodies (MVBs) that are highly acidic and degradation-prone.

As shown in Figure 2F, MC-402-103 demonstrated decreased colocalization with all examined endosomal markers compared to MC-000-101. Quantitative analysis in Figure 2G revealed that the percentage of intracellular particles associated with APPL1, EEA1, Rab11, and LBPA was reduced from 13% to 7%, 26% to 11.9%, 32.3% to 9.5%, and 32% to 14%, respectively, for MC-402-103 relative to MC-000-101. These findings support the superior endosomal escape capacity of MC-402-103, indicating that MC-402-103 was less likely to undergo recycling or degradation following internalization, leading to enhanced intracellular delivery, cytosolic release, and gene expression, as shown in Figure 1.

### Intracellular Dynamics of Particles and Lysosomes Revealed Higher Recycling Activity of MC-000-101 Compared to MC-402-103

It has been shown that endosomal escape becomes increasingly difficult as endosomes mature into lysosomes due to the more acidic environment (∼pH 5.5)^25, 26^, where cargo is more likely to be degraded before it can be released. Therefore, we further investigated and compared lysosomal interaction with LNP and pLNP particles via live-cell imaging using a spinning disk PerkinElmer confocal microscope to track the intracellular movement of particles and lysosomes over a 45-minute period.

We monitored intracellular lysosomes labeled with LysoTracker Green and DiI-labeled particles at three distinct time intervals: 30–35 min, 35–40 min, and 40–45 min (Figure 3). Matlab analysis indicated that in the MC-000-101 group, both lysosomes and particles exhibited decreasing mobility over time (from 30 to 45 min) (Figure 3A, B). At 30–35 minutes, lysosomal trafficking was prominent, suggesting active endocytosis and that endosomes containing particles had matured into lysosomes (Figure 3A). Following this, the mobility of both lysosomes and particles declined (35–40 min) and became nearly static at 40–45 min. The reduction in both lysosomal and particle trajectories implied the loss of particles, likely due to degradation or recycling. Notably, as intracellular particle trajectories declined, extracellular trajectories increased—indicating that internalized MC-000-101 particles were recycled back to the extracellular medium during this time (Figure 3B).

**Figure 3.**
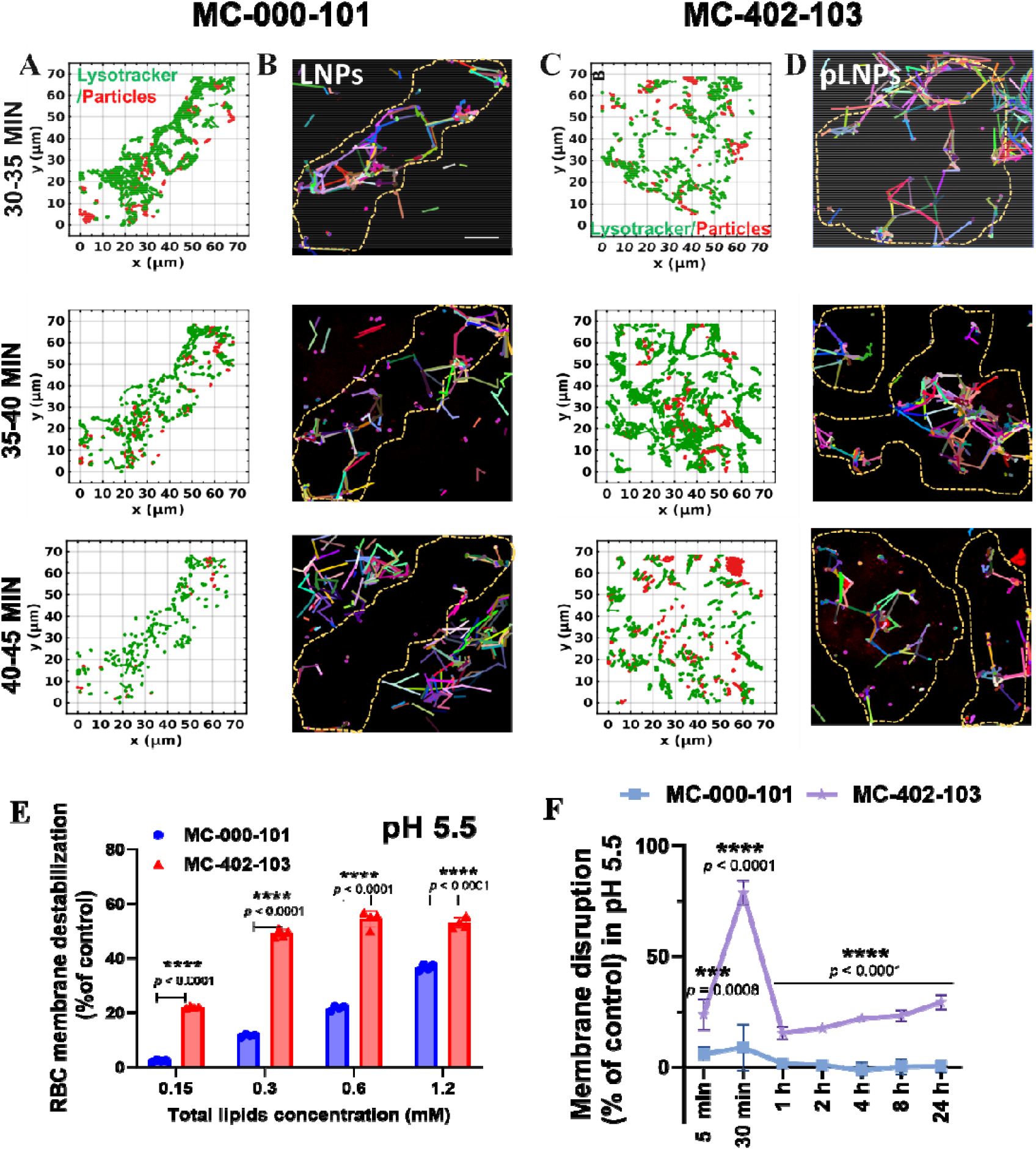
Dynamic interaction between particles and lysosomes. HEK293 cells were seeded in a confocal dish and were imagined immediately by SDCM after particles were added to the culture medium. Dynamic analyses of lysosomes (LysoTracker) and DiI-labeled particles at 30-35, 35-40, and 40-45 min after cells were incubated with MC-000-101 (**A** and **B**) and MC-402-103 (**C** and **D**) using Matlab and Fiji. **A** and **C** panels show interactions between lysosomes (green) and particles (red). **B** and **D** panels show the trails of particles (yellow dash line indicates cell edges). scale bar = 20 µm. Data was presented as lysosomes and particles trajectories over time. **E**. Red blood cell (RBC) membrane destabilization (% control) of MC-000-101 and MC-402-103 at pH 5.5. **F**. Comparison of membrane disruption after treatment with MC-000-101 or MC-402-103 at pH 5.5. FRET characterization after mixing with anionic endosomal mimics for 24 hours. Statistical significances determined using two-tailed unpaired t-test (95% CI). Data are presented as mean ± SD (n=3).

In contrast, the mobility of lysosomes and MC-402-103 particles was low during 30– 35 minutes, increased at 35–40 minutes, and then decreased again at 40–45 minutes (Figure 3C and Suppl. Figure 5A). This pattern suggests that lysosomal maturation of the endosomes enclosing MC-402-103 occurred predominantly at 35–40 minutes, approximately 5 minutes later than that of MC-000-101. This might be due to that the majority of MC-402-103 were not trapped in the lysosomes, and thus the overall lysosomal trajectories were not active at 30-35 min. Starting at 35 minutes, accumulation of the particles in endosomes that matured to lysosomes became pronounced, leading to increased lysosomal activity. At 40-45 minutes, the mobility of lysosomes decreased over time as MC-402-103 released. At 35-45 minutes, while MC-402-103 particles showed decreasing mobility overtime, their moving velocity within the cells was still higher than that of MC-000-101, suggesting increased MC-402-103 release from the lysosomes (Suppl. Figure 5B). The area of colocalization between lysosomes and MC-402-103 decreased overtime also supports the lysosomal release (Suppl. Figure 5C). Interestingly, very few MC-402-103 particles were detected outside the cells, suggesting decreased recycling activity compared to MC-000-101 (Figure 3D).

### Mechanistic Studies Demonstrate pLNPs Possesses a Superior Endosomal Membrane Destabilization Ability

We hypothesized that MC-402-103 enhanced membrane-disruptive activity to facilitate endosomal escape. The increased pH buffering effect in Suppl. Figure 4 suggests that MC-402-103 would be more effectively protonated during the acidification within endosomes compared to MC-000-101, making MC-402-103 carry more positive charges to mediate increased interaction with the anionic endosomal membrane for cytosolic release. To test this hypothesis, the red blood cell (RBC) membrane destabilization assay was conducted. While neither formulation caused disruption at neutral pH (Suppl. Figure 6A), MC-402-103 induced strong, dose-dependent membrane destabilization at pH 5.5, reaching 55% efficiency at 0.3 mM compared to just 10% for MC-000-101 (Figure 4E).

**Figure 4.**
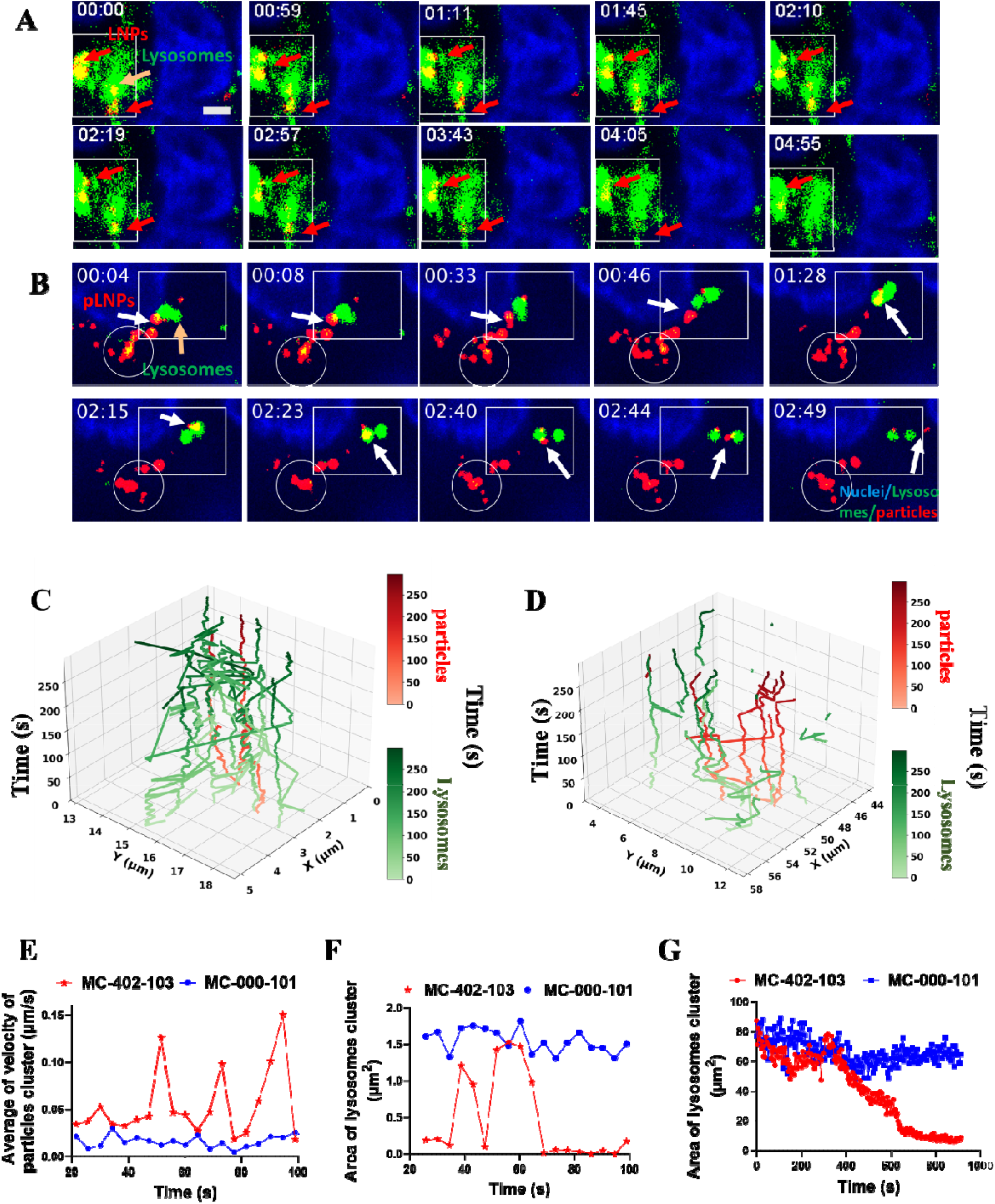
Interactions between LNP/pLNP particles and lysosomes. **A**. Squares show LNP (MC-000-101) being trapped in lysosomes. **B**. Squares show kiss- and-run: pLNP (MC-402-103) and lysosome moved directly toward each other and, upon collision, departed in opposite directions. Circles highlight lysosomal disruption mediated by pLNP, respectively. The images are arranged in chronological order. Blue: nuclei; green: lysosome (indicated by yellow arrow), red: LNP or pLNP (indicated by red and white arrow, respectively). Scale bar = 0.5 μm. Trails of particles (red) and lysosomes (green) in cells analyzed by Matlab after incubation with MC-000-101 (**C**) and MC-402-103 (**D**) from videos 1 and 2. **E**. Representative average velocity of particle clusters within cells at different time points after incubation in videos 1 and 2 in 80s. **F**. Representative total area of lysosomal clusters within cells at different time points after incubation in videos 1 and 2 in 80s. **G**. Quantification of the total area occupied by lysosomal clusters from 30 to 45 minutes within the entire field of view.

To gain deeper mechanistic insights, we employed a fluorescence resonance energy transfer (FRET) assay using DOPE-conjugated probes (DOPE-NBD as donor and DOPE-Rho as acceptor) incorporated into endosome-mimicking liposomes. When membrane disruption occurs, the increased distance between the FRET pair reduces energy transfer, leading to enhanced NBD fluorescence (Figure 4F). pLNPs induced rapid membrane destabilization, reaching 23% disruption within 5 min and up to 75% at 30 minutes. The subsequent signal reduction to 20% at 1 hour reflects thermodynamic reorganization of the lipid components, where the disrupted endosomal membranes spontaneously reassemble into energetically favorable nanostructures. The membrane-disrupting activity gradually increased to 35% at 24 hours demonstrates pLNPs’ ability to maintain continuous membrane perturbation while their inactivity at physiological pH (Suppl. Figure 6B) confirms biocompatibility. Conventional LNPs (MC-000-101) showed minimal disruption (<10%) at all timepoints. The better membrane-disrupting effect of branched NS402 might be attributed to its pH-sensitive imidazole group, and the terminal blockage modification made the polymer amphiphilic, allowing the particles to acquire cubosome-like properties with fusogenic potential—enabling membrane disruption.

### MC-402-103 escaped lysosomes via lysosomal disruption and kiss-and-run mechanisms

We further looked into the interaction between lysosomes and particles by tracking their trajectories separately in cells after incubation with MC-000-101 and MC-402-103 (Suppl. Videos 1 and 2, respectively). In MC-000-101-treated cells, particles (red) were trapped within lysosomes (green) and gradually disappeared, as indicated by the white squares in Suppl. Video 1 and Figure 4A, analyzed over a 5-min duration. In contrast, Suppl. Video 2 showed that many MC-402-103 particles were not associated with lysosomes, suggesting efficient release. Circles in Figure 4B appeared to capture the lysosomal disruption event: At 00.04, lysosomes were associated with MC-402-103 particles, emerging as significant yellow dots, which then gradually disappeared over a period of 1-2 minutes. Interestingly, the released MC-402-103 particles continued interacting with lysosomes and displayed the kiss-and-run phenomenon. Figure 4B showed that released MC-402-103 particles (red) encountered lysosomes (green) at 00:05 from different trajectories and then separated at 00:33. They “kissed” again at 00:54 and the particle merged into the lysosome becoming a yellow dot at 01:28, followed by separation into distinct red and green dots again at 02:44.

We further reconstructed the intracellular trafficking observed in Suppl. Videos 1 and 2 into Figures 4C and 4D by compiling 71 frames along the Z-axis for spatial analysis. In Figure 4C, red arrows show that MC-000-101 particles (red) remained closely associated with lysosomes (green), suggesting lysosomal trapping. In contrast, Figure 4D shows that MC-402-103 particles (red) exhibited little interaction with lysosomes. High particle mobility typically indicates endo-lysosomal escape or encapsulation in early endosomal vesicles, while low mobility reflects entrapment in late endosomes or lysosomes^27^. Figure 4E confirms that MC-402-103 maintained relatively high velocity (0.025–0.15 µm/s) even during lysosomal interactions, suggesting low entrapment within lysosomes. The occasional fluctuations in velocity—periods of slowing down followed by recovery—may indicate transient lysosomal trapping. However, once the ionizable groups on MC-402-103 became protonated in the acidic lysosomal environment, they would likely destabilize the lysosomal membrane, promoting dissociation of the particles from the lysosomes. After escaping, the particles rapidly returned to their original velocity. Conversely, MC-000-101 showed a lower velocity (0.01–0.02 µm/s) once trapped in lysosomes as their moving area was constrained inside the lysosomes.

Analysis of lysosomal cluster areas in Suppl. Video 1 and 2 (Figure 4F) revealed that lysosomes in MC-000-101-treated cells maintained a consistent size (∼1.5 µm^2^) over time. In contrast, the lysosomes within the MC-402-103-treated cells displayed significant area fluctuations (0–1.5 µm^2^), consistent with proton sponge-induced swelling (1.5 µm^2^) and subsequent rupture (0.2 µm^2^), supporting the proton sponge mechanism. Furthermore, Figure 4G shows that the total lysosomal area in MC-402-103-treated cells decreased over time from 30 to 45 minutes, whereas cells treated with MC-000-101 exhibited no significant change, indicating limited lysosomal rupture.

### MC-402-103 Enabled Efficient Delivery of Adenosine Deaminase Base Editors (ABEs) for *in vivo* Genome Editing

We then compared the efficacy of MC-402-103 and MC-000-103 in delivering RNAs for genome editing in a transgenic mice model (LumA). This reporter mouse model carries the R387X mutation (c.A1159T) in the luciferase gene within the Rosa26 locus. This mutation abolishes luciferase activity, which can be restored through A-to-G correction by a SpCas9 ABEs, as illustrated in Figure 5A. We formulated ABEs mRNA and sgRNA into MC-000-101 and MC-402-103 and i.v. administered them to the LumA mice, followed by monitoring of genome editing over 14 days. As shown in Suppl. Table 6, MC-402-103 had a mean size of 74 nm with a PDI < 0.1, while MC-000-101 were 84 nm with a PDI of 0.2. Both formulations carried a slightly negative charge (∼ –2 mV) and exhibited comparable RNA EE% > 90%. As shown in Figure 5B, the gene editing in both groups became detectable by day 3 for the liver luciferase bioluminescence, which increased in intensity over time and plateaued at day 12. Notably, the MC-402-103 formulation produced an 8-fold stronger bioluminescence signal than MC-000-101, with quantification presented in Figure 5C. On day 14, mice were euthanized, and major organs were collected. Sanger sequencing of genomic DNA from mouse liver samples showed that MC-402-103 achieved an 8% T-to-C conversion, whereas MC-000-101 resulted in only ∼1% (Figure 5D). Remarkably, this high editing efficiency was achieved with a 1/20 dose (0.1 mg/kg) than typically used in similar studies (2 mg/kg)^4^. These results demonstrate the high delivery efficiency by MC-402-103.

**Figure 5.**
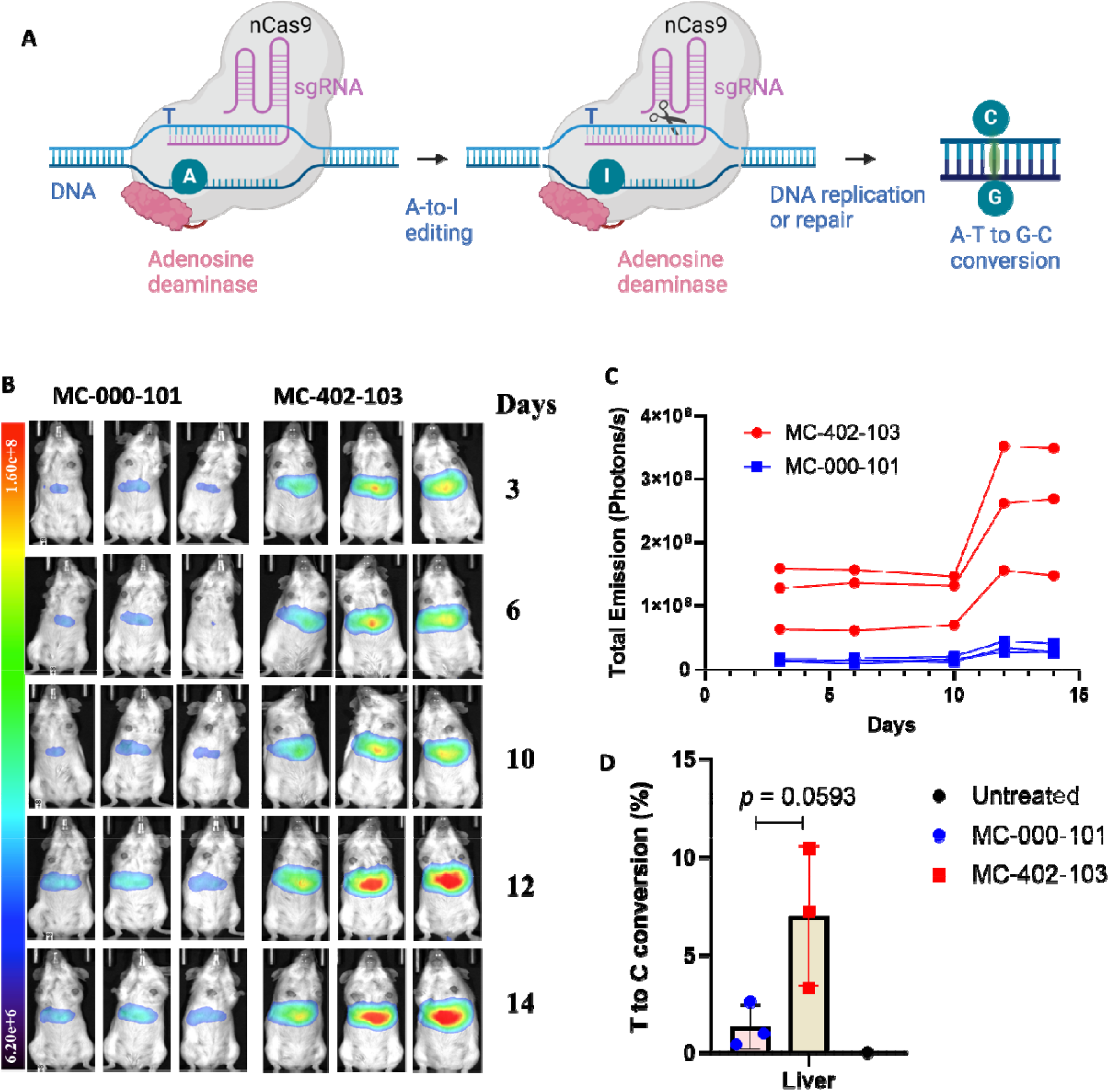
Delivery of mRNA encoded with adenosine deaminase base editor (ABE) for genome editing in LumA mice. **A**. Gene editing mechanism of ABE. Bioluminescence images (**B**) of and the quantitative results (**C**) in the LumA mic after i.v. injection of particles loaded with ABE mRNA and sgRNA (data=mean ± SD, n=3). **D**. Quantification of base editing in the liver after different treatments.

### Safety Evaluation of MC-402-103

*In vitro* cytotoxicity studies of MC-402-103 in HEK 293 cells showed no significant changes in viability, even at a high lipid concentration of 500 µM (Suppl. Figure. 7A). *In vivo* immunotoxicity was examined by analyzing the serum collected from mice 3 hours post i.v. injection of MC-402-103 at 0.3 mg/kg. As shown in Suppl. Figure. 7B and C, TNF-α and IL-6 levels show no statistical difference compared to PBS control, indicating no immunotoxicity. Given the liver’s predominant role in particle accumulation, hepatic toxicity was assessed one day post-injection through serum biomarker analysis including renal function markers (e.g., creatinine, blood urea nitrogen (BUN)); electrolytes and metabolic panel (e.g., phosphorus, total anions, osmolality); protein and liver enzymes (e.g., total protein, albumin, Alanie aminotransferase (ALT)) and interference indices (e.g., hemolysis index, icterus index). As summarized in Suppl. Table 7, all biomarker levels remained within normal physiological ranges, with no significant differences among the treatment groups, suggesting no overt liver toxicity.

### Machine Learning-Guided Optimization of Polyhistidine (H_10_) configuration to prepare enhanced pLNPs

Building upon our empirical findings that symmetrical bis-lysine histidine dendron (e.g., NS402) exhibited 2-fold higher transfection efficiency than its linear counterpart (NS310) at comparable MW and mol%, we recognized configuration as a critical but underexplored design parameter, yet the vast design space for branched H_10_ presents a critical challenge in selecting optimal structures. Conventional structure-activity approaches struggle to capture three-dimensional arrangement effects, as evidenced by the dramatic performance differences between NS402 (branched) and NS404 (branched but less effective) despite similar molecular compositions. To address this issue, we turned to ML to decode how topological features, including branching degree (linear vs. dendrimeric) and histidine spatial density, govern transfection efficiency. We aimed to both identify predictive features and guide the design of next-generation pLNPs while minimizing the need for exhaustive screening. Despite its potential, ML workflows designed for predictions in the small-molecule space can struggle to adequately describe macromolecules. This is especially notable when applying classical ML strategies with features such as binary molecular fingerprints. Most established descriptors designed for small molecules (10–100 atoms) rely on atom-level resolution and well-defined bonding patterns. However, when applied to large, flexible structures like polyhistidines, these descriptors become impractical— failing to capture branching patterns and sequence-dependent effects that dictate function. Instead of an atomistic approach, we deployed an amino-acid level graph representation, where nodes correspond to amino acids or protecting group functionalities and edges represent peptide bonds. By condensing structural complexity into a simplified yet chemically relevant framework, this strategy enabled ML models to detect meaningful structure–function relationships, particularly in small-data environments like the one presented here.

To implement an ML-guided optimization strategy, we compiled experimental data from Figure 1, incorporating polyhistidine structures, peptide mol%, and bioluminescence intensities at various time points. Each polyhistidine peptide was featured as described above and fed into a GNN with three graph convolution layers based on the GNN operator from Morris et al^24^ (Figure 6A and B). Given the large scale discrepancy and noise in data measurements, separate models were trained on the minimum, maximum, and average ln(maximum radiance) across all runs (Figure 6A). The final model provided high prediction statistics with R^2^ = 0.86, train mean squared Error (MSE) = 0.53, Q^2^ = 0.87, and test MSE = 0.36 on a random 80:20 train test split (Figure 6B). Furthermore, good predictions were observed in a leave-one-peptide-out test, a scaffold-split test where each peptide was removed from training, the model retrained, and the excluded peptide data predicted. Despite no representation in the training set, most peptides were predicted reasonably well with 10/24 with MSE within 1, 15/24 with MSE within 2, and 20/24 with MSE within 3, demonstrating that this model should reasonably extrapolate to new structural regimes. Notably, this model greatly outperformed benchmark models built using atom-wise graph representations or molecular fingerprints (Suppl. machine learning logics), further supporting the validity of the amino-acid level featurization.

**Figure 6.**
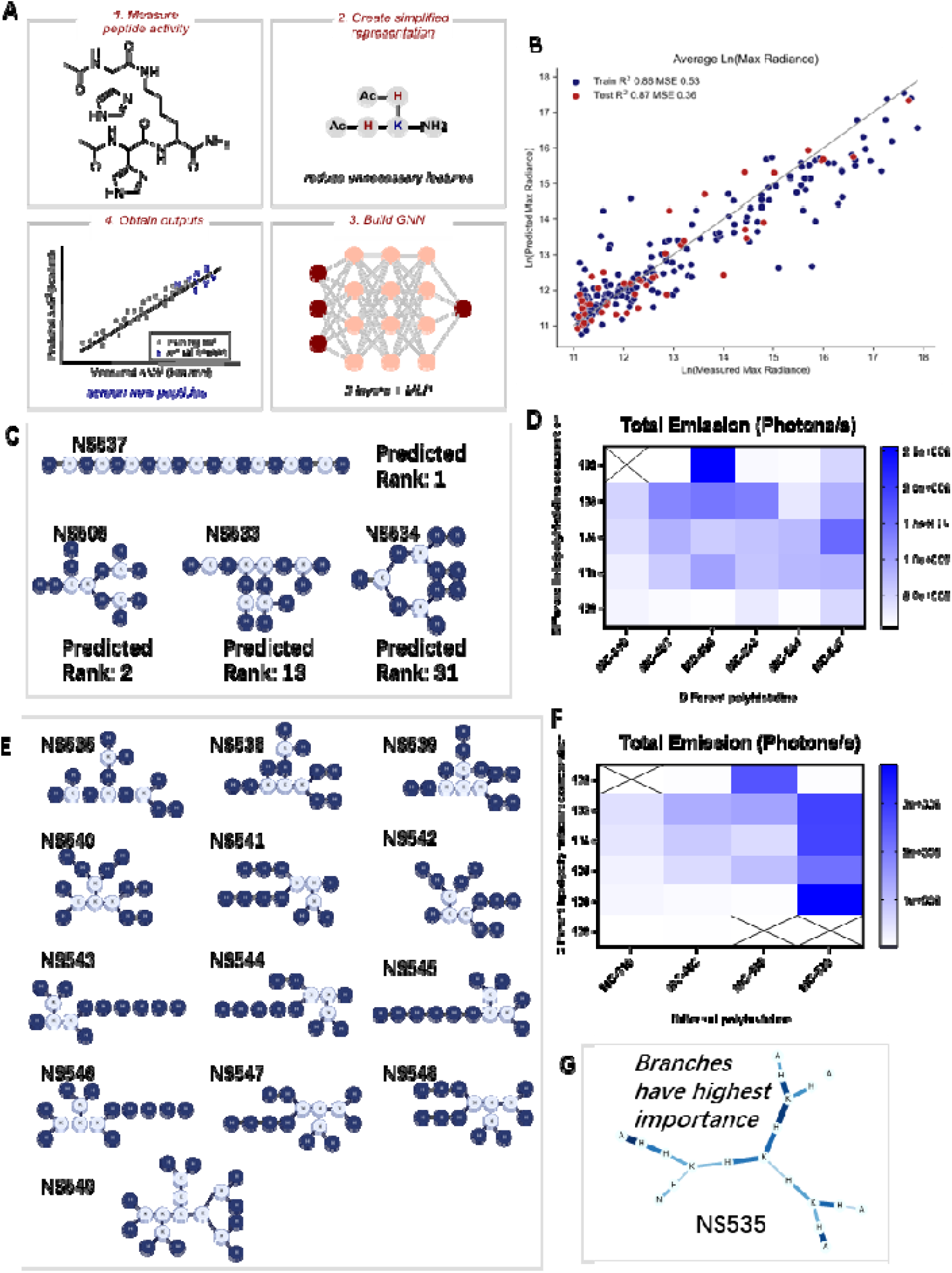
Machine learning–guided optimization and validation of polyhistidine (H_10_) conformation for enhanced *in vivo* mRNA transfection. **A**. Work flow of the ML model. MLP: multilayer perceptron; GNN: Graph Neural Network. First, pooled data from Figure 1B and G, Suppl. Figure 2 and Suppl. Figure 3B. Second, created graph representation where amino acids are nodes, and not atoms. Third, built GNN to predict bioluminescence from peptide structures. Fourth, used the trained GNN to predict and validate the model. **B**. New peptide structures were predicted by ML, which indicated branched structures were more effective compared to linear. **C**. Structures of NS506, NS533 and NS537 suggested by the ML method that would improve the transfection efficiency. NS534 was predicted to be inferior to NS402. **D**. Total flux of bioluminescence in the liver 5 h after i.v. delivery of different pLNP formulations loaded with luciferase mRNA (n=3). X-axis represents different polyhistidines (refer to Figure. 6C for their structures) used to prepare pLNP formulations. Y-axis represents different lipid/polyhistidine compositions (refer to Suppl. Table 1 for compositions). **E**. Structures of newly designed branched H_10_ for ML model prediction. **F**. Total flux of bioluminescence in the liver 5 h after i.v. delivery of different pLNP formulations loaded with luciferase mRNA (n=3). X-axis represents different polyhistidines (refer to Figure. 6E for their structures) used to prepare pLNP formulations. Y-axis represents different lipid/polyhistidine compositions (refer to Suppl. Table 1 for compositions). **G**. Structural model of NS535. Wider and darker edges corresponding to higher importance.

Using this model, we prospectively designed polyhistidine sequences (NS501–534 and NS537; Suppl. machine learning logics: virtual screening round 1). Notably, model predictions (Figure 6C) ranking NS506 and NS537 highest for *in vivo* testing while NS533 and NS534 were ranked lower (13^th^ and 31^st^, respectively) (Suppl. Excel Sheet 1). As shown in Supplementary Table 8, pLNPs incorporating NS506 and NS537 exhibited average diameters slightly above 100 nm (PDI <0.3), while NS533 and NS534 formed smaller nanoparticles (∼90 nm, PDI <0.15 for NS533, <0.3 for NS534), with encapsulation efficiencies (EE%) >70% (Suppl. Table 8). *In vivo*, NS506 demonstrated a 2-fold increase in luciferase bioluminescence compared to NS402, while NS533 and NS537 enhanced expression 1.9-fold and 1.1-fold respectively, confirming the predictive ability of the model (Figure 6D).

Incorporating the newly gathered bioluminescence data from NS506, NS533, NS534, and NS537, we refined the model and performed a second round of virtual screening (Suppl. machine learning logics: virtual screening round 2) with designed peptides NS535, NS538–NS549, prioritizing the star-like branched H_10_ scaffold (Figure 6E). Here, sequences NS535 and NS545 ranked the highest amongst the screened peptides (Suppl. Excel Sheet 2), and NS535 was selected as a high-performing candidate to test *in vivo*. NS535-containing pLNPs exhibited comparable sizes (∼100 nm, PDI <0.25) and EE% (Suppl. Table 9). Importantly, MC-535-108 showed a 2.2-fold increase in liver bioluminescence over MC-506-102 and a 2.9-fold increase over MC-402-103, validating our ML-driven design strategy (Figure 6F). This final optimized formulation achieved a 705-fold increase in mRNA expression compared to the initial standard formulation (MC-000-101) (3.1*10^9^ vs. 4.4*10^6^ Photons/s, *p* = 0.0031).

Having demonstrated that ML can guide the optimization of polyhistidine peptide structure, we last questioned if important structural features could be parsed using explainable ML techniques. A new GNN was trained using the same procedure as described above but without the concatenation of additional graph-level features. This purely structural model was then used to predict NS535, where histidine residues radiate symmetrically from a central lysine core, and more importantly, served as a proxy to uncover why NS535 was predicted to lead to such high transfection efficiency. The GNNExplainer by Ying et al^28^ was employed to explain important subgraphs in this model and the results shown in Figure 6G. Here, the sum of edge importance is denoted by the shade and width of the edge, with wider and darker edges corresponding to higher importance. As expected, the highest importance is given to the branched portions of the structure, further suggesting that the branched dendritic configuration with histidine residues densely arranged on a multi-valent core is necessary for high transfection.

This approach accelerated the screening of macromolecular conformations by ranking 48 polyhistidine configurations, providing a predictive model for optimizing nanoparticle formulations and quantifying how structural modifications influenced biological performance. By implementing an innovative amino acid-level graph representation, we overcame the limitations of traditional small-molecule descriptors, enabling accurate prediction of polyhistidine architectures that optimized mRNA transfection. This computational strategy is uniquely suited for macromolecular systems, capturing the intricate structure-function relationships that drive delivery efficiency.

## Conclusion

In this study, we established a systematic strategy to engineer pLNPs for enhanced mRNA delivery by elucidating the SAR of polyhistidine architectures. Our findings showed that branched polyhistidine designs—particularly those based on H_10_ scaffolds—significantly improved transfection efficiency, achieving up to a 266-fold increase compared to conventional LNPs. Mechanistic studies revealed that these optimized pLNPs efficiently escaped the endosomal network, promoting mRNA delivery through two synergistic mechanisms: proton sponge-induced endosomal swelling and disruption, and kiss-and-run dissociation from lysosomes. We delivered ABE mRNA using pLNPs and achieved 8% editing efficiency in the liver of mice at a remarkably low dose, underscoring their clinical relevance. Finally, we developed a ML framework based on amino acid–level graph representations. This model accurately predicted high-performing polyhistidine structures and identified branching topology as a key driver of delivery efficiency. The top candidate, NS535, promoted a 705-fold enhancement in hepatic gene transfection relative to standard LNPs. Our work established polyhistidine configuration - alongside MW and mol% - as a fundamental design parameter for LNP delivery systems. The demonstration that spatial arrangement determined delivery efficiency (with branched configurations outperforming linear analogs by 2-3 fold) represents a paradigm shift in polymer engineering for nucleic acid delivery. The ML framework’s ability to decode these configuration-activity relationships provides a blueprint for rational design of next-generation delivery systems. Together, this work presents an integrated platform that combines rational design, mechanistic insight, therapeutic validation, and ML-guided optimization to advance mRNA delivery technologies. By uniting data-driven prediction with in vivo functional testing, our approach offers both foundational understanding of polymeric SAR and a scalable strategy for developing next-generation nucleic acid carriers.

## Supporting information

Supplemental information 2-machine learning logics

Supplemental excel sheet 1

Supplemental excel sheet 2

