## Supplemental information 1-tables and figures for "Machine Learning–Guided Structure–Activity Discovery of Polymer Configurations in Lipid Nanoparticles for Kiss-and-Run Endosomal Escape"

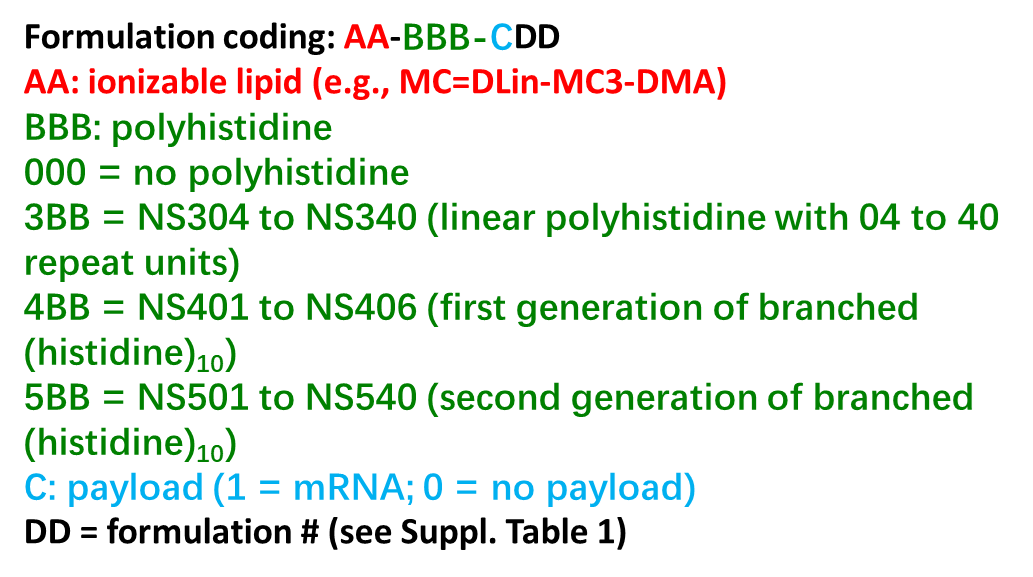

**Suppl. Figure 1.** The formulation coding logics (mRNA = luciferase mRNA)

**Suppl. Table 1:** Mole percentages of each component in LNP and pLNP formulations

| **Formulation #** | **01** | **02** | **03** | **04** | **05** | **08** | **09** |
| --- | --- | --- | --- | --- | --- | --- | --- |
| **MC3** | 50% | 48.78% | 47.62% | 45.5% | 41.67% | 33.3% | 25.0% |
| **DSPC** | 10% | 10% | 10% | 10% | 10% | 10% | 10% |
| **Cholesterol** | 38.5% | 38.5% | 38.5% | 38.5% | 38.5% | 38.5% | 38.5% |
| **DMG-PEG** | 1.5% | 1.5% | 1.5% | 1.5% | 1.5% | 1.5% | 1.5% |
| **Peptide** | 0% | 1.22% | 2.38% | 4.5% | 8.33% | 16.7% | 25.0% |

**Suppl. Table 2:** Size, PDI, Zeta potential and encapsulation efficiency (EE %) of LNP and pLNP formulations.

| **Name** | **Size (nm)** | **PDI** | **Zeta Potential** | **EE%** |
| --- | --- | --- | --- | --- |
| MC-000-101 | 70.91±1.43 | 0.15±0.04 | -1.76±1.23 | 93.7 |
| MC-304-103 | 78.21±7.84 | 0.20±0.02 | -3.09±2.74 | 70.2 |
| MC-304-104 | 85.44±17.04 | 0.18±0.04 | 0.25±0.38 | 75.3 |
| MC-304-105 | 78.19±9.16 | 0.17±0.05 | -1.18±0.51 | 74.4 |
| MC-304-108 | 79.56±9.62 | 0.23±0.03 | -1.17±1.12 | 58.4 |
| MC-304-109 | 73.40±7.66 | 0.20±0.05 | -0.22±0.09 | 70.2 |
| MC-305-103 | 82.89±19.78 | 0.13±0.01 | -9.51±8.08 | 69.4 |
| MC-305-104 | 69.79±0.97 | 0.10±0.04 | -5.16±5.49 | 72.4 |
| MC-305-105 | 77.73±16.59 | 0.20±0.06 | -6.77±4.00 | 55.5 |
| MC-305-108 | 62.31±1.10 | 0.12±0.01 | -17.70±1.08 | 64.4 |
| MC-305-109 | 84.85±2.06 | 0.01±0.01 | -2.19±1.18 | 18.6 |
| MC-306-103 | 70.17±1.07 | 0.12±0.01 | -1.50±1.04 | 65.4 |
| MC-306-104 | 85.21±1.80 | 0.12±0.03 | -0.70±0.60 | 65.9 |
| MC-306-105 | 81.14±5.74 | 0.14±0.01 | -0.91±0.34 | 68.0 |
| MC-306-108 | 82.71±4.17 | 0.11±0.01 | 0.22±1.51 | 61.7 |
| MC-306-109 | 77.72±2.74 | 0.17±0.01 | -0.91±1.19 | 56.2 |
| MC-307-103 | 74.80±1.33 | 0.13±0.01 | -4.67±6.56 | 76.8 |
| MC-307-104 | 79.83±9.05 | 0.20±0.05 | -0.68±0.62 | 70.5 |
| MC-307-105 | 74.89±3.56 | 0.19±0.01 | -2.16±2.80 | 78.6 |
| MC-307-108 | 102.53±1.14 | 0.25±0.01 | -2.71±0.59 | 92.8 |
| MC-307-109 | 81.99±6.92 | 0.27±0.02 | -0.34±0.47 | 20.5 |
| MC-308-103 | 59.93±1.84 | 0.15±0.03 | -2.29±0.94 | 63.8 |
| MC-308-104 | 85.93±2.43 | 0.12±0.02 | -0.76±0.51 | 63.4 |
| MC-308-105 | 73.64±18.56 | 0.21±0.03 | -0.60±0.79 | 69.7 |
| MC-308-108 | 71.09±13.28 | 0.20±0.01 | -1.92±0.53 | 54.7 |
| MC-308-109 | 84.74±2.13 | 0.22±0.01 | -1.90±1.02 | 73.0 |
| MC-309-103 | 74.62±0.90 | 0.16±0.00 | -1.08±0.35 | 85.7 |
| MC-309-104 | 72.08±0.69 | 0.14±0.03 | -0.44±0.05 | 88.0 |
| MC-309-105 | 68.77±2.26 | 0.15±0.04 | -0.02±0.36 | 88.5 |
| MC-309-108 | 67.75±1.44 | 0.15±0.01 | -1.19±0.35 | 89.4 |
| MC-309-109 | 77.13±0.99 | 0.12±0.02 | -8.75±0.72 | 80.6 |
| MC-310-103 | 79.70±0.25 | 0.14±0.03 | -0.12±0.15 | 83.8 |
| MC-310-104 | 76.45±0.25 | 0.09±0.02 | -9.89±1.30 | 91.4 |
| MC-310-105 | 67.55±1.05 | 0.09±0.01 | -0.12±1.28 | 90.7 |
| MC-310-108 | 79.41±0.79 | 0.15±0.01 | -5.29±1.18 | 74.2 |
| MC-310-109 | 72.23±0.84 | 0.13±0.01 | 0.17±0.15 | 41.5 |
| MC-311-103 | 86.26±1.36 | 0.06±0.01 | 2.31±0.19 | 65.9 |
| MC-311-104 | 81.32±0.91 | 0.07±0.01 | -0.05±0.10 | 90.8 |
| MC-311-105 | 80.59±1.34 | 0.11±0.03 | 0.60±0.30 | 91.7 |
| MC-311-108 | 72.68±0.56 | 0.13±0.01 | 0.18±0.37 | 54.2 |
| MC-311-109 | 74.74±1.01 | 0.21±0.01 | -0.50±0.62 | 47.0 |
| MC-312-103 | 90.90±6.56 | 0.12±0.02 | -0.12±0.34 | 62.7 |
| MC-312-104 | 97.68±0.46 | 0.15±0.01 | -2.83±0.12 | 84.0 |
| MC-312-105 | 88.00±1.53 | 0.17±0.01 | -2.03±0.30 | 78.1 |
| MC-312-108 | 80.22±1.38 | 0.17±0.01 | -1.80±2.14 | 66.0 |
| MC-312-109 | 73.87±0.54 | 0.21±0.02 | -0.02±0.38 | 46.1 |
| MC-313-103 | 106.07±3.00 | 0.17±0.03 | -0.07±0.12 | 83.0 |
| MC-313-104 | 98.97±1.93 | 0.18±0.01 | 0.03±0.21 | 79.7 |
| MC-313-105 | 90.73±1.59 | 0.16±0.01 | -0.39±0.27 | 72.1 |
| MC-313-108 | 106.50±2.04 | 0.12±0.02 | -0.19±0.19 | 56.3 |
| MC-313-109 | 244.77±21.54 | 0.24±0.01 | -0.85±0.50 | 68.1 |
| MC-314-103 | 90.19±1.21 | 0.14±0.00 | -0.13±0.33 | 78.3 |
| MC-314-104 | 71.57±0.84 | 0.11±0.01 | -0.10±0.25 | 63.4 |
| MC-314-105 | 78.55±2.31 | 0.21±0.01 | 0.73±1.04 | 52.8 |
| MC-314-108 | 92.44±12.46 | 0.25±0.03 | -1.43±1.05 | 82.6 |
| MC-314-109 | 127.87±5.85 | 0.22±0.03 | -1.18±0.22 | 93.2 |
| MC-315-103 | 100.65±1.52 | 0.18±0.01 | -0.25±0.29 | 57.3 |
| MC-315-104 | 93.46±1.95 | 0.24±0.01 | 2.61±1.06 | 78.5 |
| MC-315-105 | 110.57±1.19 | 0.25±0.01 | -5.83±0.57 | 44.5 |
| MC-315-108 | 653.47±550.68 | 0.69±0.31 | -2.06±0.72 | 23.5 |
| MC-315-109 | 281.03±69.91 | 0.37±0.20 | -1.37±0.80 | 22.4 |
| MC-316-103 | 99.95±1.46 | 0.15±0.02 | -2.85±1.16 | 92.8 |
| MC-316-104 | 127.17±2.78 | 0.14±0.04 | -0.25±0.09 | 73.0 |
| MC-316-105 | 159.37±58.56 | 0.24±0.02 | -0.27±0.10 | 23.9 |
| MC-316-108 | 136.60±1.42 | 0.24±0.03 | -0.35±0.20 | NA |
| MC-316-109 | 1,038.43±775.89 | 0.74±0.26 | -0.36±0.11 | NA |
| MC-324-102 | 137.17±0.79 | 0.13±0.03 | -1.94±0.51 | 93.0 |
| MC-324-103 | 550±57.1 | 0.26±0.04 | 2.39±2.39 | 93.8 |
| MC-324-104 | 162.55±11.89 | 0.24±0.06 | -4.91±9.01 | 85.1 |
| MC-324-105 | 11,683.00±15,482.93 | 0.79±0.24 | -1.62±2.74 | 80.6 |
| MC-324-108 | 21,819.00±12,860.81 | 0.85±0.17 | 4.39±8.19 | 78.1 |
| MC-332-102 | 126.53±10.22 | 0.26±0.06 | -0.91±1.15 | 88.3 |
| MC-332-103 | NA | 0.01±0.01 | -0.92±0.93 | 55.7 |
| MC-332-104 | 4,240.67±5,255.01 | 0.95±0.05 | -0.34±2.06 | 41.9 |
| MC-332-105 | 719±68.8 | 0.28±0.04 | 8.3±0.15 | 83.7 |
| MC-340-102 | 436.60±409.95 | 0.41±0.36 | -0.41±0.07 | 78.6 |
| MC-340-103 | 6,238.00±3,352.10 | 1.00±0.00 | 0.66±2.89 | 16.4 |
| MC-340-104 | 594.8±74.9 | 0.25±0.01 | 5.51±1.07 | 52.2 |
| MC-340-105 | 771.67±1,373.75 | 1.00±0.00 | No Data | 9.9 |

**Suppl. Table 3:** Size, PDI, Zeta potential and encapsulation efficiency (EE %) of pLNP formulations.

| **Name** | **Size (nm)** | **PDI** | **Zeta Potential** | **EE%** |
| --- | --- | --- | --- | --- |
| MC-310-103 | 79.70±0.25 | 0.14±0.03 | -0.12±0.15 | 83.9 |
| MC-310-104 | 76.45±0.25 | 0.09±0.02 | -9.89±1.30 | 91.4 |
| MC-310-105 | 67.55±1.05 | 0.09±0.01 | -0.12±1.28 | 90.7 |
| MC-310-108 | 79.41±0.79 | 0.15±0.01 | -5.29±1.18 | 74.2 |
| MC-310-109 | 72.23±0.84 | 0.13±0.01 | 0.17±0.15 | 41.5 |
| MC-310n-103 | 84.17±0.76 | 0.07±0.02 | 0.63±0.17 | 37.6 |
| MC-310n-104 | 84.73±0.97 | 0.13±0.01 | 0.96±0.63 | 35.8 |
| MC-310n-105 | 97.01±1.54 | 0.11±0.00 | 0.54±0.31 | 50.11 |
| MC-310n-108 | 117.77±3.11 | 0.10±0.00 | 0.46±0.02 | 51.2 |
| MC-310n-109 | 183.5±5.63 | 0.24±0.02 | -0.17±0.49 | 56.4 |

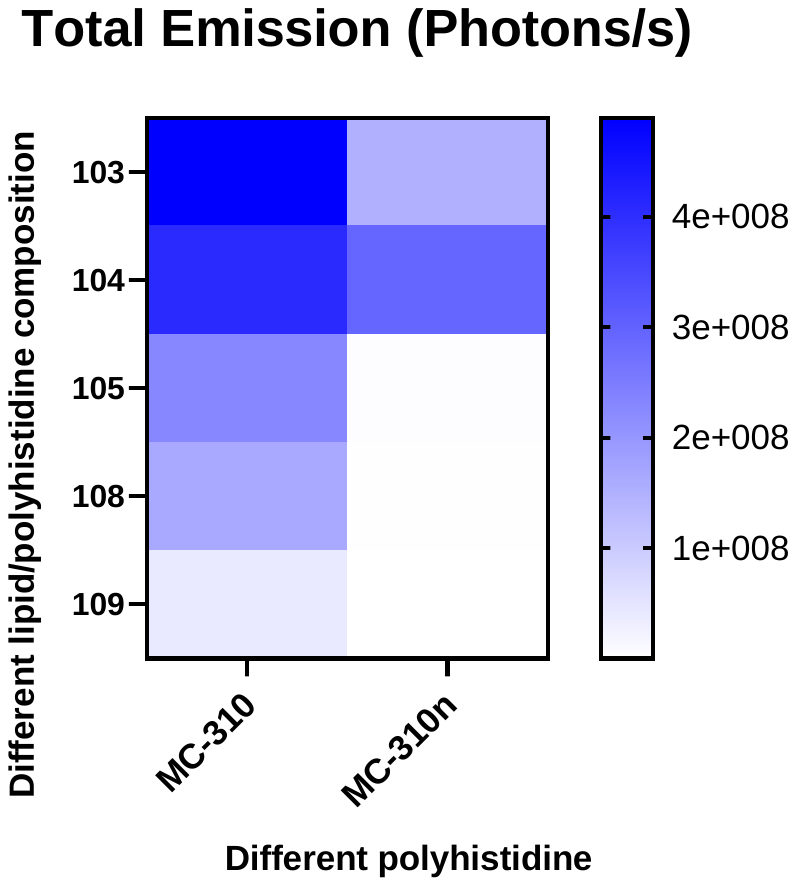

**Suppl. Figure 2.** Total flux of bioluminescence in the liver 5 h after I.V. delivery of different pLNP formulations loaded with luciferase mRNA (n=3). X-axis represents different polyhistidines (refer to Suppl. Figure. 1 for their structures) used to prepare pLNP formulations. MC-310n represents that the N and C termini in the H_10_ peptide were not blocked. Y-axis represents different lipid/polyhistidine compositions (refer to Suppl. Table 1 for compositions).

**Suppl. Table 4:** Size, PDI, Zeta potential and encapsulation efficiency (EE %) of pLNP formulations.

| **Name** | **Size (nm)** | **PDI** | **Zeta Potential** | **EE%** |
| --- | --- | --- | --- | --- |
| MC-401-103 | 77.88±2.03 | 0.12±0.01 | -0.94±0.84 | 73.9 |
| MC-401-104 | 83.72±1.95 | 0.10±0.02 | -0.12±1.13 | 85.4 |
| MC-401-105 | 90.40±4.85 | 0.14±0.02 | -0.10±0.26 | 80.2 |
| MC-401-108 | 94.42±1.28 | 0.20±0.01 | -0.26±1.18 | 49.7 |
| MC-401-109 | 92.56±1.16 | 0.22±0.01 | 0.91±1.48 | 19.1 |
| MC-402-103 | 91.13±0.90 | 0.11±0.02 | 0.30±0.61 | 90.1 |
| MC-402-104 | 76.12±0.47 | 0.10±0.01 | -12.03±2.74 | 81.5 |
| MC-402-105 | 83.45±1.75 | 0.12±0.01 | -7.25±1.88 | 76.6 |
| MC-402-108 | 81.09±1.52 | 0.21±0.01 | -2.55±1.28 | 52.7 |
| MC-402-109 | 115.97±1.88 | 0.16±0.01 | -3.28±0.27 | 58.6 |
| MC-403-103 | 85.03±1.25 | 0.09±0.01 | -0.69±1.05 | 78.2 |
| MC-403-104 | 77.22±1.23 | 0.13±0.01 | -8.88±2.76 | 70.8 |
| MC-403-105 | 75.72±5.13 | 0.17±0.06 | -8.14±3.01 | 85.9 |
| MC-403-108 | 94.96±2.51 | 0.28±0.02 | 0.84±0.23 | 50.8 |
| MC-403-109 | 81.56±1.72 | 0.29±0.03 | 1.43±0.13 | 55.8 |
| MC-404-103 | 79.03±6.01 | 0.20±0.02 | 1.52±1.03 | 61.0 |
| MC-404-104 | 86.03±1.57 | 0.06±0.03 | 1.29±0.31 | 89.8 |
| MC-404-105 | 78.69±0.50 | 0.19±0.02 | 0.85±0.95 | 51.4 |
| MC-404-108 | 88.60±2.17 | 0.13±0.01 | 0.97±0.53 | 57.2 |
| MC-404-109 | 93.21±13.15 | 0.28±0.07 | 0.65±0.46 | 29.7 |

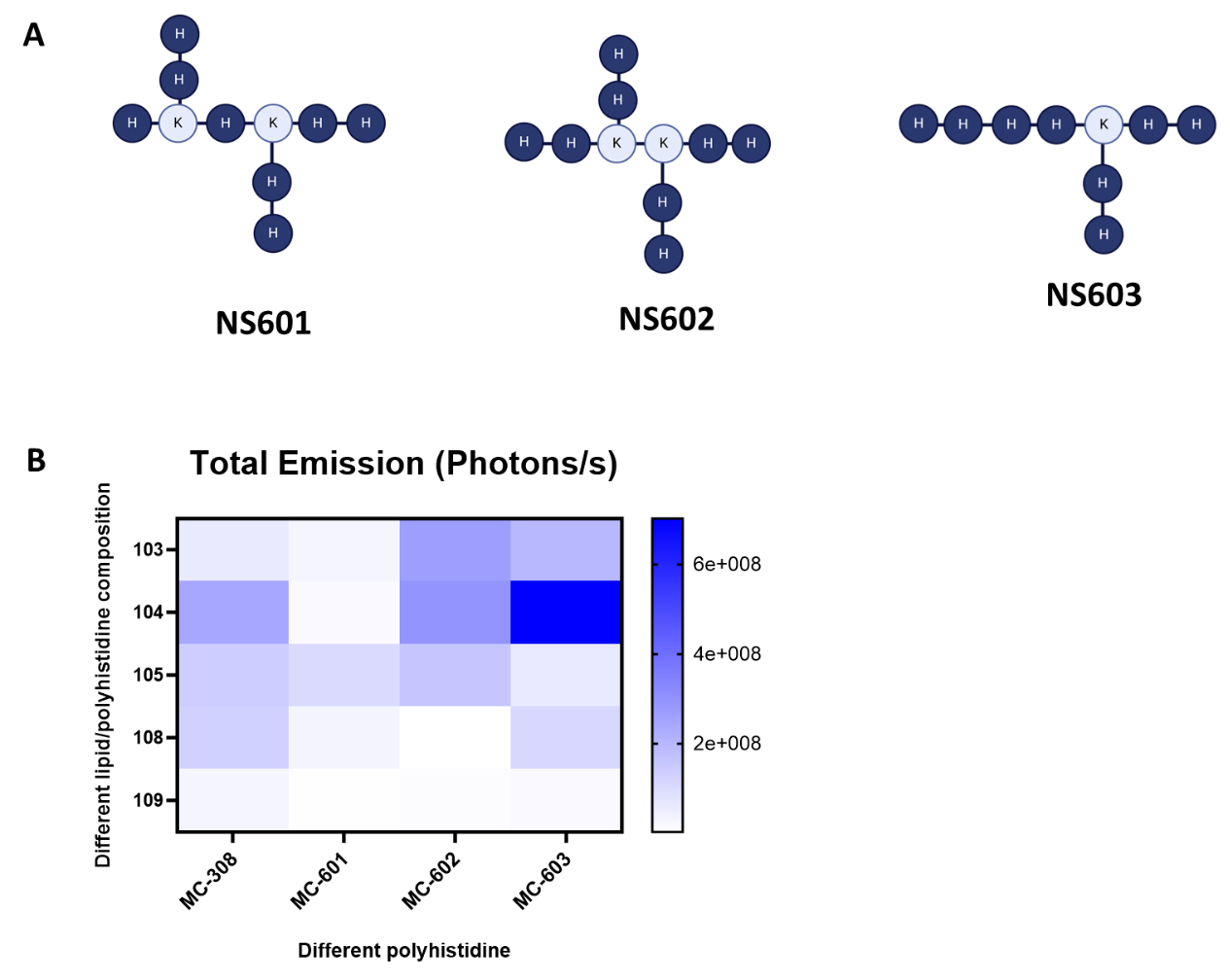

**Suppl. Figure 3.** pLNP formulations prepared with different branched H_8_. **A**. structures of branched H_8_. **B**. Total flux of bioluminescence in the liver 5 h after i.v. delivery of different LNP and pLNP formulations loaded with luciferase mRNA (n=3). X-axis represents different polyhistidines (different numbers of repeat units of histidine) used to prepare pLNP formulations. Y-axis represents different lipid/polyhistidine compositions. (refer to Suppl. Table 1 for compositions).

**Suppl. Table 5:** Size, PDI, Zeta potential and encapsulation efficiency (EE %) of pLNP formulations.

| **Name** | **Size (nm)** | **PDI** | **Zeta Potential** | **EE%** |
| --- | --- | --- | --- | --- |
| MC-601-103 | 87.71±3.70 | 0.13±0.02 | -0.06±0.13 | 91.6 |
| MC-601-104 | 82.90±0.59 | 0.14±0.01 | -0.69±0.96 | 89.7 |
| MC-601-105 | 86.89±1.14 | 0.20±0.01 | -1.69±0.41 | 72.2 |
| MC-601-108 | 82.20±2.14 | 0.15±0.01 | 0.92±0.92 | 73.8 |
| MC-601-109 | 86.25±1.73 | 0.14±0.01 | -1.74±1.80 | 82.9 |
| MC-602-103 | 71.48±2.37 | 0.13±0.02 | -0.36±1.43 | 72.2 |
| MC-602-104 | 89.64±32.21 | 0.22±0.02 | -0.80±0.79 | 90.0 |
| MC-602-105 | 69.38±5.26 | 0.17±0.05 | -0.42±0.70 | 89.5 |
| MC-602-108 | 57.82±0.48 | 0.16±0.03 | -0.61±0.26 | 86.8 |
| MC-602-109 | 64.46±1.57 | 0.17±0.01 | -1.31±1.16 | 84.2 |
| MC-603-103 | 81.51±1.11 | 0.21±0.01 | -4.12±1.22 | 84.9 |
| MC-603-104 | 63.35±1.40 | 0.11±0.01 | -2.32±2.34 | 86.4 |
| MC-603-105 | 74.51±2.45 | 0.21±0.01 | -6.68±11.69 | 89.8 |
| MC-603-108 | 64.12±3.84 | 0.15±0.05 | -1.77±0.81 | 87.4 |
| MC-603-109 | 76.02±2.98 | 0.12±0.04 | -13.00±9.02 | 87.0 |

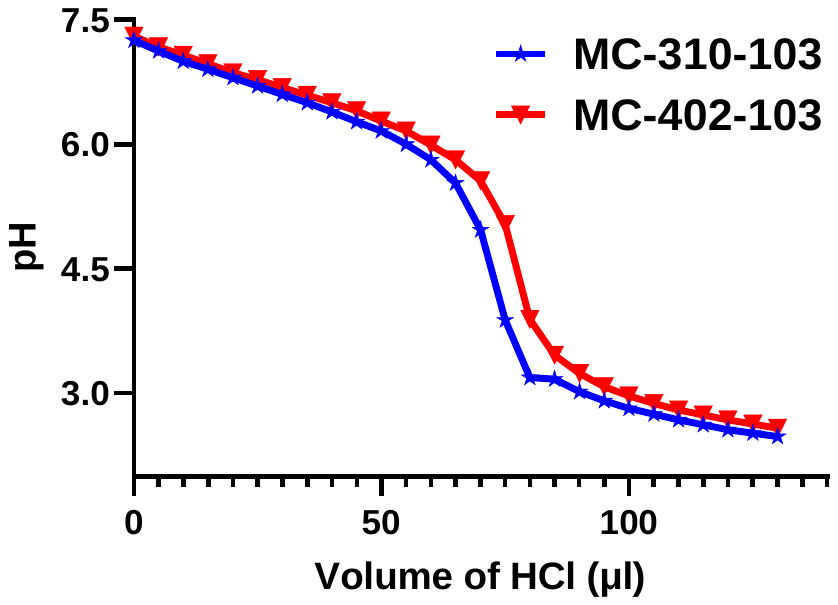

**Suppl. Figure 4**. pH buffering effect of linear and branched polyhistidine incorporated pLNP formulations measured by the acid titration assay.

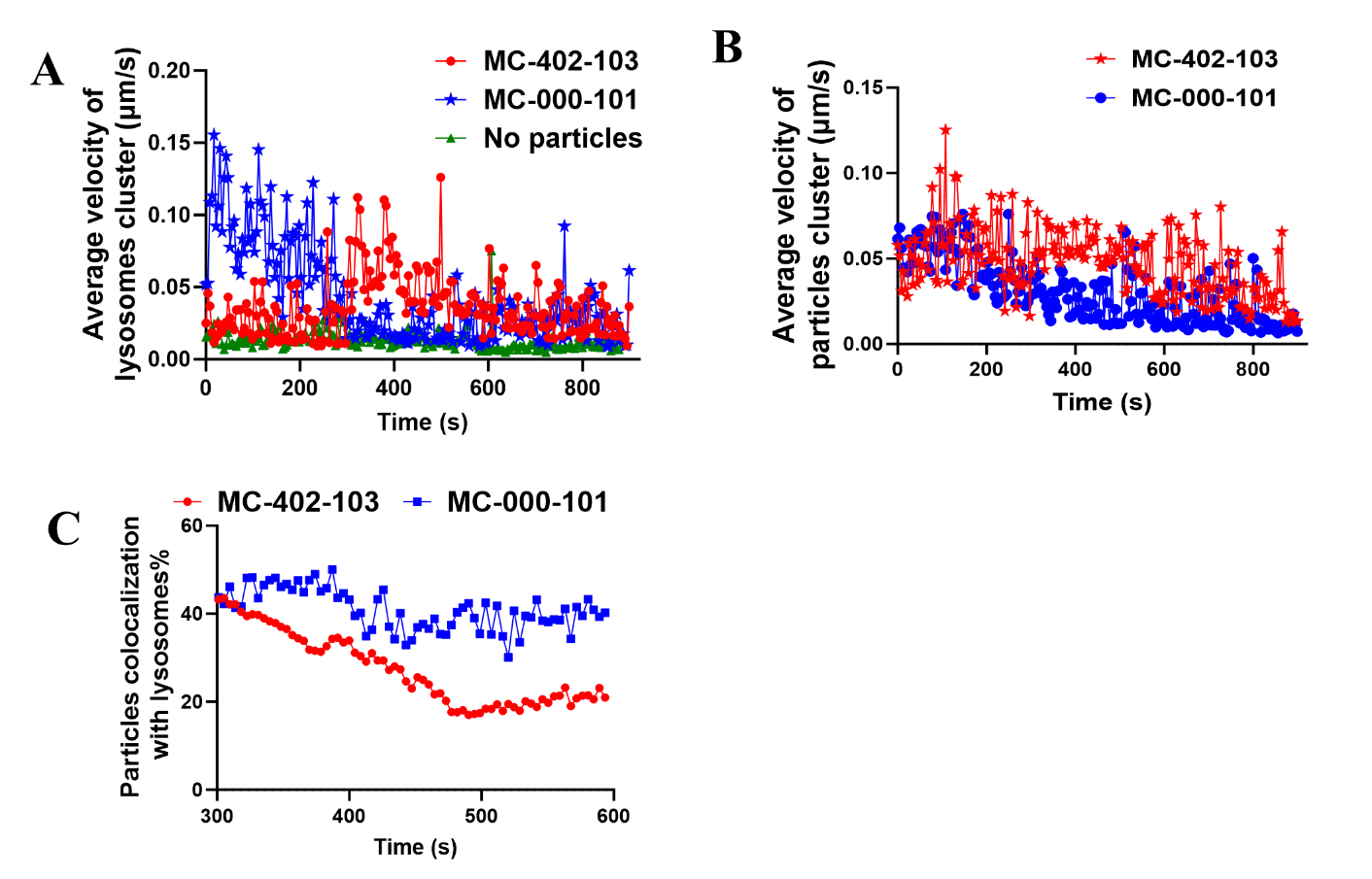

**Suppl. Figure 5. Dynamic interaction of particles with lysosomes.** Analyzing videos of cells after treatment with MC-000-101 or MC-402-103 at 30 to 45 min by using Fiji to show the velocity of endosomes (**A**), and particles (**B**). Percentage of particles and lysosomes overlapping from 35 to 40 min(**C**).

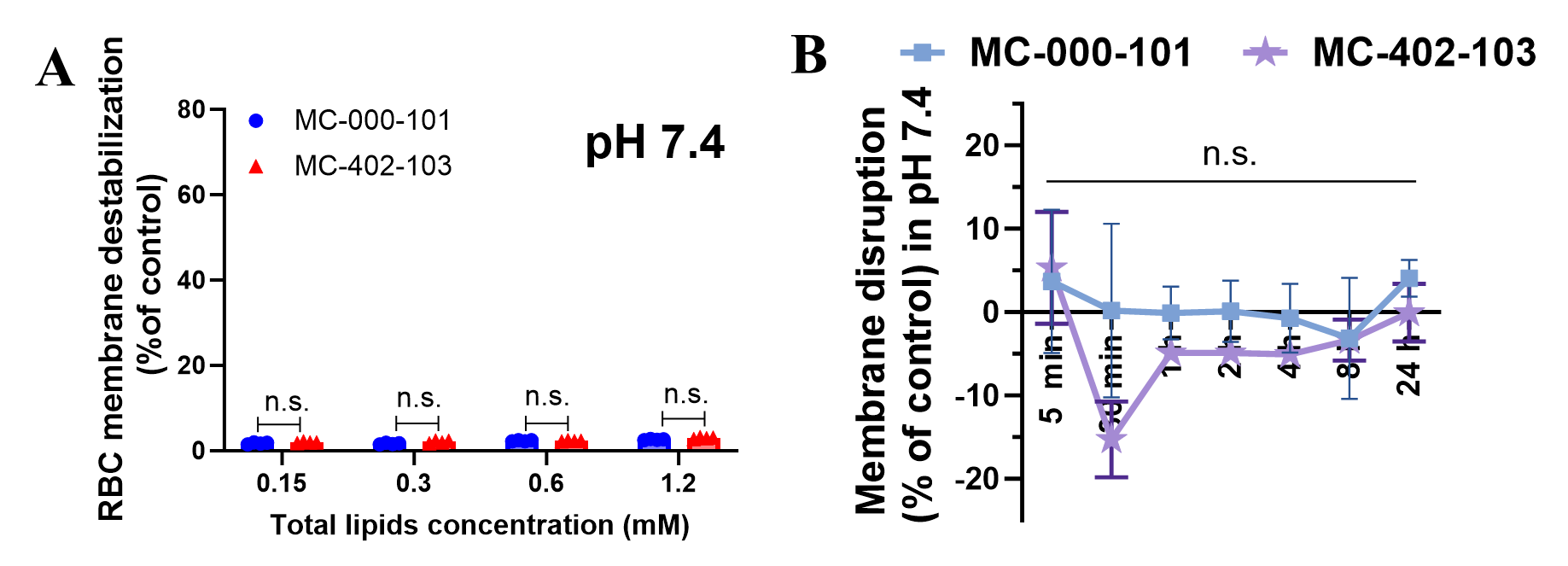

**Suppl. Figure 6. Mechanistic studies of particle mediated endosomal escape.** Red blood cells (RBC) membrane destabilization (% control) of MC-000-101 and MC-402-103 at pH 7.4 (**A**). Comparison of membrane disruption with MC-000-101 and MC-402-103 at 7.4. by FRET characterization after mixing with anionic endosomal mimics for 24 hours (**B**). Statistical significances determined using two-tailed unpaired *t*-test (95% CI). Data are presented as mean ± SD, n.s. no significant difference (n=3).

**Suppl. Table 6:** Size, PDI, Zeta potential and encapsulation efficiency (EE %) of LNP and LNP formulations containing ABE mRNA and sgRNA

| **Name** | **Size (nm)** | **PDI** | **Zeta Potential** | **EE%** |
| --- | --- | --- | --- | --- |
| MC-402-103 | 74.3±0.90 | 0.08±0.07 | -2.08±0.33 | 92.2 |
| MC-000-101 | 83.4±5.7 | 0.20±0.02 | -1.78±1.07 | 95.0 |

**Suppl. Table7:** Safety evaluation function 24 h after mice receive 0.3 mg/kg pLNPs via i.v.

|  |  |  | |  | **MC-402-103** | **PBS** |
| --- | --- | --- | --- | --- | --- | --- |
|  |  | **Unit** | | **Normal Range** | **Mean±SD** | **Mean±SD** |
| **Creatinine** |  | mg/dL | | 0.2-0.4 | 0.1±0 | <0.1 |
| **Urea (BUN)** | | mg/dL | 15-59 | | 23.0±1.7 | 20.1±3.6 |
| **BUN: Creatinine Ratio** | |  |  | | 201.7±81.9 | 200.5±35.3 |
| **Phosphorus** |  | mg/dL | | 6.0-11.3 | 8.4±0.8 | 7.0±1.5 |
| **Sodium** |  | mmol/L | | 145-175 | 147.7±1.9 | 152.8±1.3 |
| **Potassium** |  | mmol/L | | 6.5-9.7 | 4.7±1.0 | >10 |
| **Calcium** |  | mg/dL | | 6.8-11.9 | 8.4±3.1 | 0±0 |
| **Chloride** |  | mmol/L | | 111-134 | 109.0±1.6 | 104.0±4.7 |
| **TCO2 (Bicarbonate)** | | mmol/L |  | | 16.3±1.6 | 9.3±1.5 |
| **Anion Gap** |  |  | | 8.8-30.8 | 27.4±2.2 | 49.5±4.5 |
| **Total Cations** | | mmol/L |  | | 152.3±1.9 | 162.8±1.3 |
| **Total Anions** | | mmol/L |  | | 125.0±2.3 | 113.3±5.7 |
| **Total Protein** | | g/dL | 3.3-6.4 | | 4.9±0.2 | 5.2±0.3 |
| **Albumin** |  | g/dL | | 2.5-3.5 | 2.8±0.2 | 3.0±0.2 |
| **Globulin** |  | g/L | |  | 20.8±1.6 | 21.5±1.3 |
| **Albumin: Globulin Ratio** | |  |  | | 1.4±0.2 | 1.4±0.1 |
| **ALT** |  | U/L | | 7-227 | 99.8±77.9 | 26.5±4.8 |
| **AST** |  | U/L | | 57-329 | 206.2±187.8 | 75.3±21.4 |
| **Bilirubin - Total** | | µmol/L | 0.1-0.9 | | 0.1±0 | 0.2±0 |
| **Osmolality** |  | mmol/kg | |  | 305.3±3.9 | 320.3±3.9 |
| **Hemolysis Index** | |  |  | | Normal | Normal |
| **Icterus Index** | |  |  | | Normal | Normal |
| **Lipemia Index** | |  |  | | Normal | Normal |

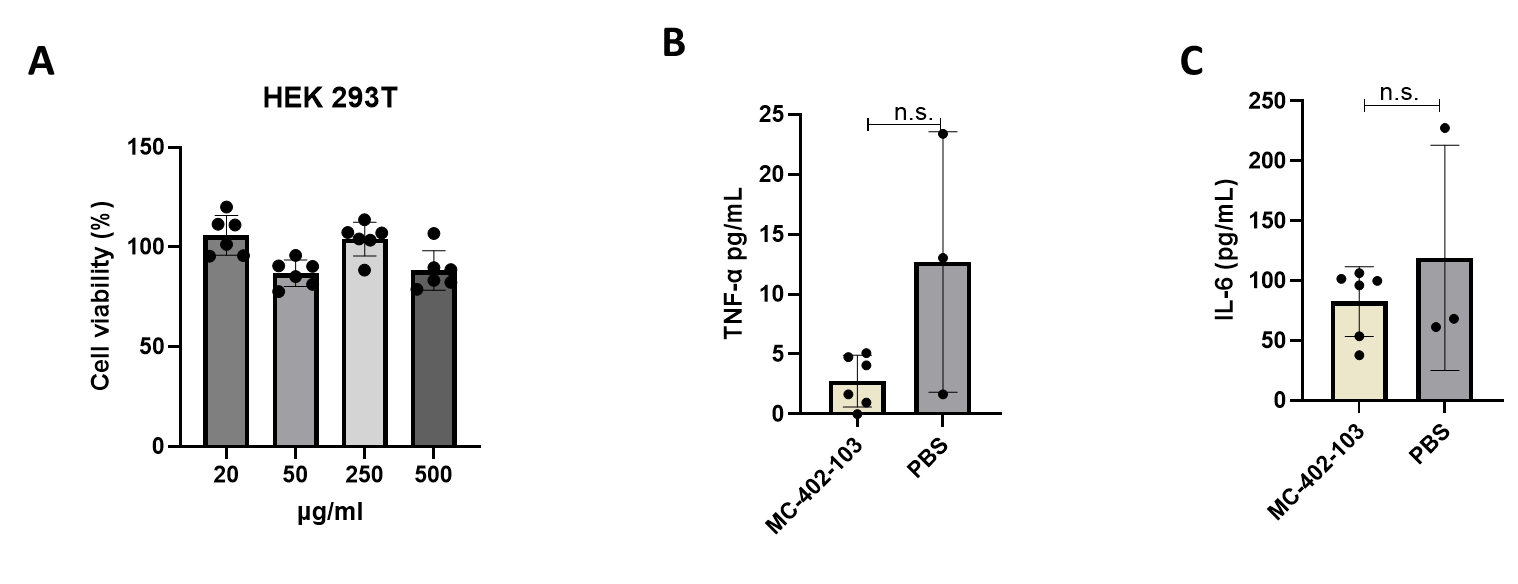

**Suppl. Figure 7. Safety evaluation of pLNPs in vitro and in vivo. A**. Cell viability of pLNPs. **B** and **C**. TNF-α and IL-6 levels in serum 3 h after mice receive pLNPs at dose 0.3 mg/kg.

**Suppl. Table 8:** Size, PDI, Zeta potential and encapsulation efficiency (EE %) of pLNP

| **Name** | **Size (nm)** | **PDI** | **Zeta Potential** | **EE%** |
| --- | --- | --- | --- | --- |
| MC-506-102 | 103.2±14.6 | 0.23±0.03 | -7.55±1.24 | 76.5 |
| MC-506-103 | 103.83±1.16 | 0.09±0.01 | -0.53±0.65 | 78.6 |
| MC-506-104 | 146.9±5.1 | 0.33±0.02 | -7.77±0.76 | 78.8 |
| MC-506-105 | 113.10±2.07 | 0.13±0.03 | -2.27±0.56 | 71.1 |
| MC-506-108 | 111.47±1.19 | 0.09±0.01 | -1.73±0.63 | 60.5 |
| MC-533-102 | 122±1.38 | 0.16±0.02 | -2.99±0.52 | 78.0 |
| MC-533-103 | 82.35±1.29 | 0.06±0.01 | -0.71±0.20 | 80.6 |
| MC-533-104 | 75.52±1.50 | 0.08±0.03 | -4.98±1.33 | 83.2 |
| MC-533-105 | 78.51±1.63 | 0.12±0.01 | -14.00±9.53 | 78.4 |
| MC-533-108 | 89.42±1.79 | 0.10±0.00 | -4.62±1.37 | 63.1 |
| MC-534-102 | 92.36±5.46 | 0.09±0.03 | -3.65±4.94 | 106.2 |
| MC-534-103 | 106.9±3.4 | 0.27±0.02 | -7.93±0.52 | 104.6 |
| MC-534-104 | 85.77±0.92 | 0.09±0.02 | -1.27±1.95 | 100.9 |
| MC-534-105 | 119.10±4.75 | 0.24±0.02 | -4.09±5.21 | 74.7 |
| MC-534-108 | 85.58±4.50 | 0.14±0.05 | -1.39±1.35 | 101.7 |
| MC-537-102 | 125.70±1.67 | 0.24±0.01 | -1.98±0.55 | 69.3 |
| MC-537-103 | 103.60±1.60 | 0.16±0.01 | -2.09±0.11 | 82.9 |
| MC-537-104 | 95.52±1.59 | 0.12±0.01 | -2.61±0.83 | 78.1 |
| MC-537-105 | 99.6±3.3 | 0.25±0.02 | -6.66±0.31 | 82.6 |
| MC-537-108 | 141.27±9.24 | 0.34±0.05 | -2.41±0.71 | 67.0 |

**Suppl. Table 9:** Size, PDI, Zeta potential and encapsulation efficiency (EE %) of pLNP

| **Name** | **Size (nm)** | **PDI** | **Zeta Potential** | **EE%** |
| --- | --- | --- | --- | --- |
| MC-535-102 | 103.1±3.3 | 0.25±0.01 | -7.4±0.6 | 85.1 |
| MC-535-103 | 95.51±0.54 | 0.07±0.04 | -13.31±11.77 | 98.2 |
| MC-535-104 | 100.02±0.66 | 0.12±0.02 | 0.00±0.03 | 92.7 |
| MC-535-105 | 97.30±3.15 | 0.09±0.04 | -0.08±0.29 | 74.1 |
| MC-535-108 | 105.57±2.40 | 0.05±0.05 | -1.03±0.56 | 62.4 |
