## Supplemental information 2-machine learning logics for "Machine Learning–Guided Structure–Activity Discovery of Polymer Configurations in Lipid Nanoparticles for Kiss-and-Run Endosomal Escape"

All code was written in python version 3.12.4. GNN models were developed using PyTorch (2.2.2) and PyTorch geometric (2.5.3). GNNExplainer was deployed using the torch. Explainer class in PyTorch geometric. Prior to model evaluation, data was randomly split into 80:20 training:validation splits. Benchmarks to classical ML algorithms were performed using scikit-learn (1.5.1).^1^ Molecular fingerprints were generated using RDKit (2024.3.5).^2^

**All code and data are available in the accompanying repository.**

**Training**

The model was trained using the Adam optimizer^4^ to minimize the mean-squared-error loss:

$$L\left( \hat{y},y \right)=\frac{1}{N}\sum_{i=1}^{N} \left( \hat{y_{i}}-y_{i} \right)^{2}$$

Hyperparameters (batch size, hidden channels, learning rate) were tuned using a grid search on the mean ln(max rad) dataset. Each run was trained for 10 000 epochs and evaluated by the test set RMSE.

**Table S1.** Hyperparameter tuning.

| Batch Size | Hidden  Channels | LR | Min test  RMSE | min test  epoch |
| --- | --- | --- | --- | --- |
| **32** | **512** | **0.01** | **0.83** | **6848** |
| 32 | 128 | 0.01 | 0.85 | 1768 |
| 32 | 256 | 0.01 | 0.87 | 6800 |
| 32 | 512 | 0.001 | 0.89 | 7278 |
| 16 | 256 | 0.01 | 0.89 | 9063 |
| 32 | 256 | 0.001 | 0.89 | 6101 |
| 32 | 128 | 0.001 | 0.92 | 4992 |
| 16 | 128 | 0.01 | 0.92 | 6857 |
| 16 | 128 | 0.001 | 0.94 | 6667 |
| 16 | 256 | 0.001 | 0.94 | 6170 |
| 16 | 512 | 0.01 | 0.95 | 8728 |
| 32 | 128 | 0.1 | 0.95 | 6721 |
| 16 | 512 | 0.001 | 0.96 | 2729 |
| 32 | 512 | 0.1 | 1.45 | 3188 |
| 32 | 256 | 0.1 | 1.48 | 2257 |
| 16 | 256 | 0.1 | 1.56 | 843 |
| 16 | 128 | 0.1 | 1.58 | 159 |
| 16 | 512 | 0.1 | 1.59 | 8455 |

Given the large discrepancy in the scale of the Max Rad values, directly learning these values proved to be difficult. We found that a simple output transformation to the ln(max radiance) provided better fits and model predictions, likely due to a more even data distribution.

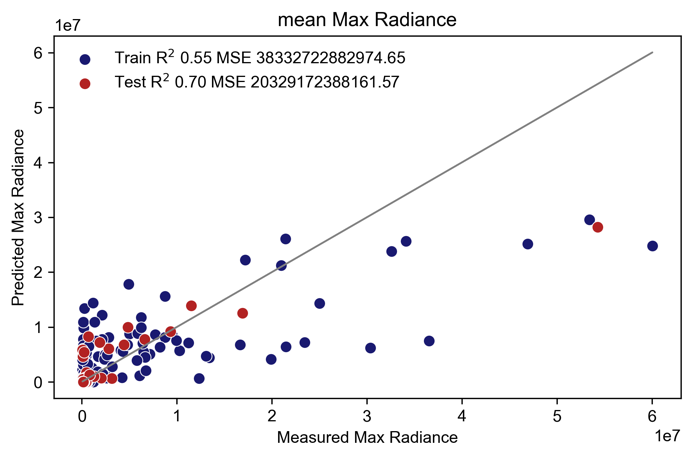

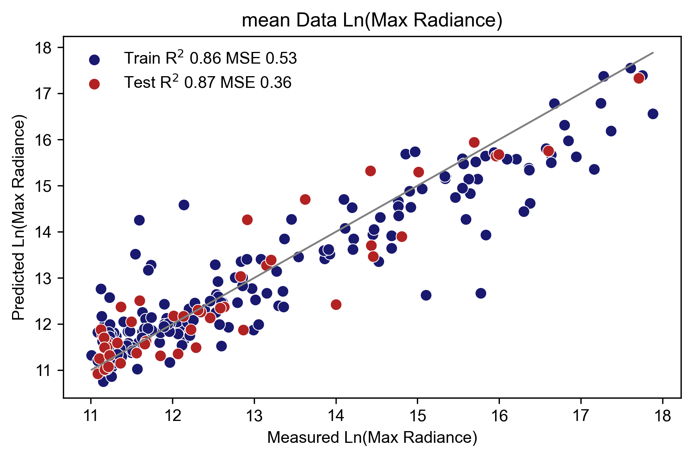

**Figure S1.** Comparison of a GNN trained with max radiance (top) or ln(max radiance) (bottom).

Furthermore, there is inherent noise with each measurement. We find that separately modelling the minimum, mean, and maximum ln(max radiance) resulted in improved model performance compared to an alternative where all data is available to the model.

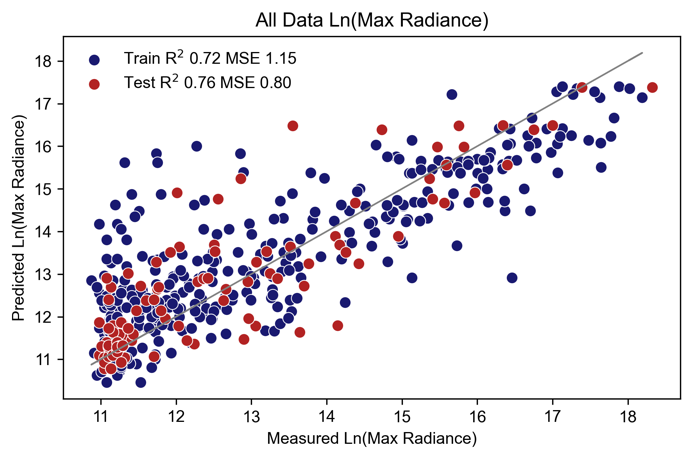

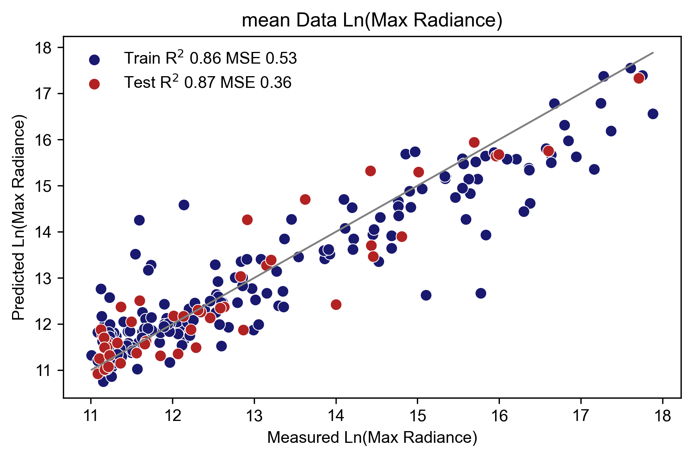

**Figure S2.** Comparison of a GNN trained on all data vs the mean of runs.

Compared to a simple linear transformation, the addition of a final MLP regressor improves model performance.

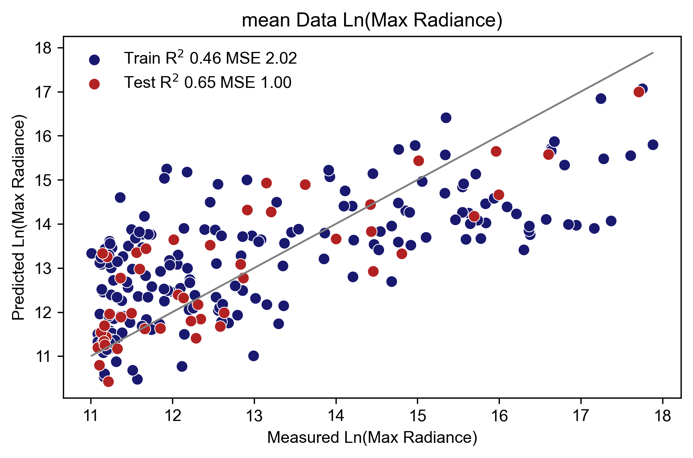

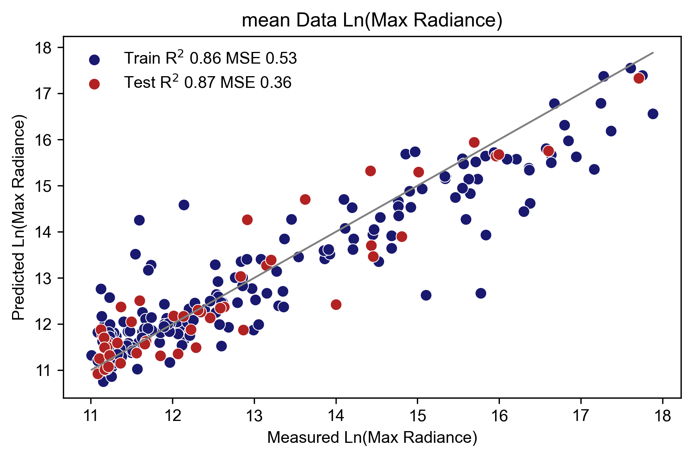

**Figure S3.** Comparison of a GNN with a final linear transformation (top) or MLP regressor (bottom).

To validate that the model is generalizable, we performed leave-one-peptide-out cross-validation. Here, each peptide is removed from the training set and predicted using the remaining peptides, simulating a realistic prediction scenario. Despite no representation in the training set, most peptides are predicted reasonably well with 10/24 with MSE within 1, 15/24 with MSE within 2, and 20/24 with MSE within 3.

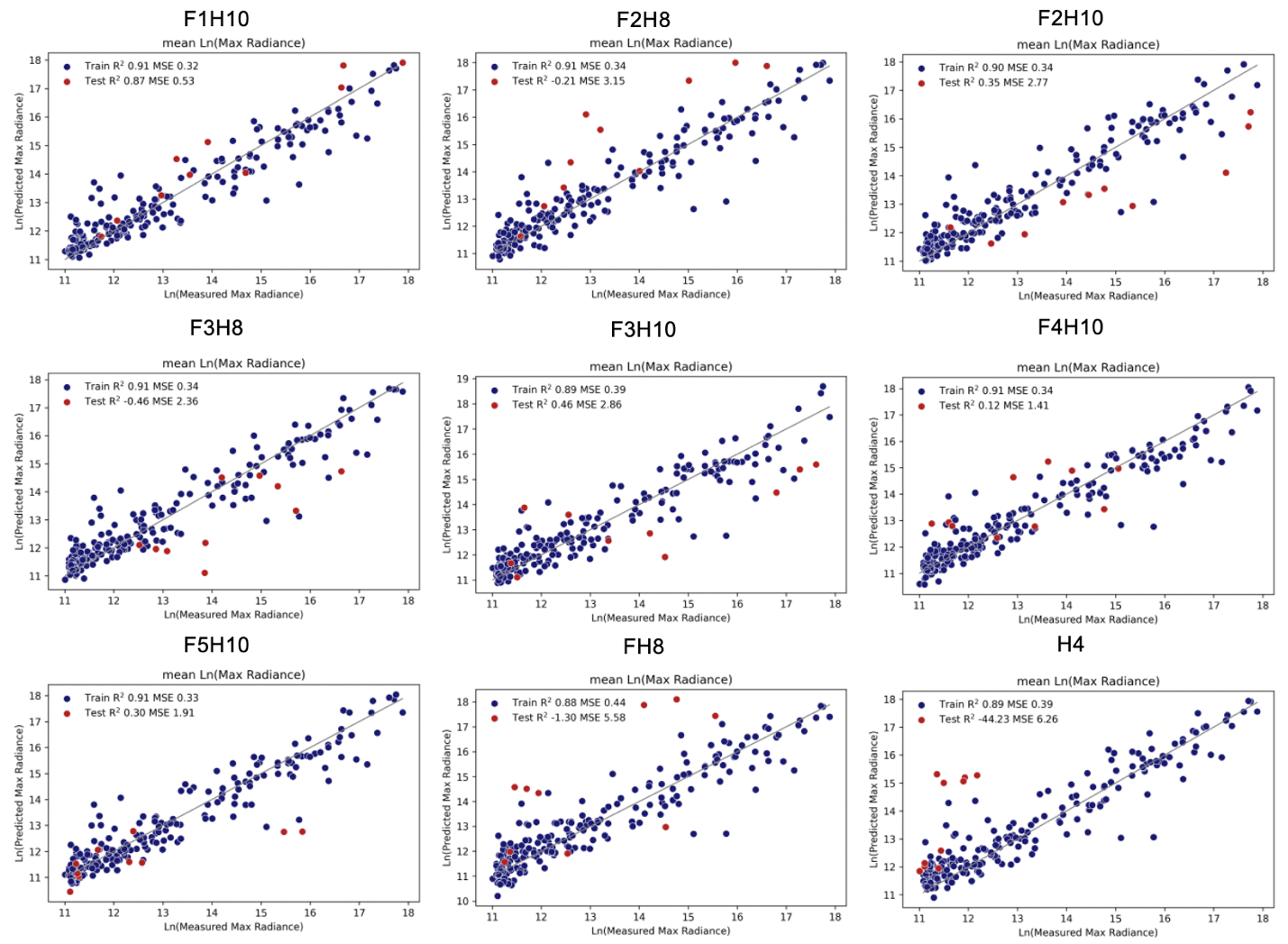

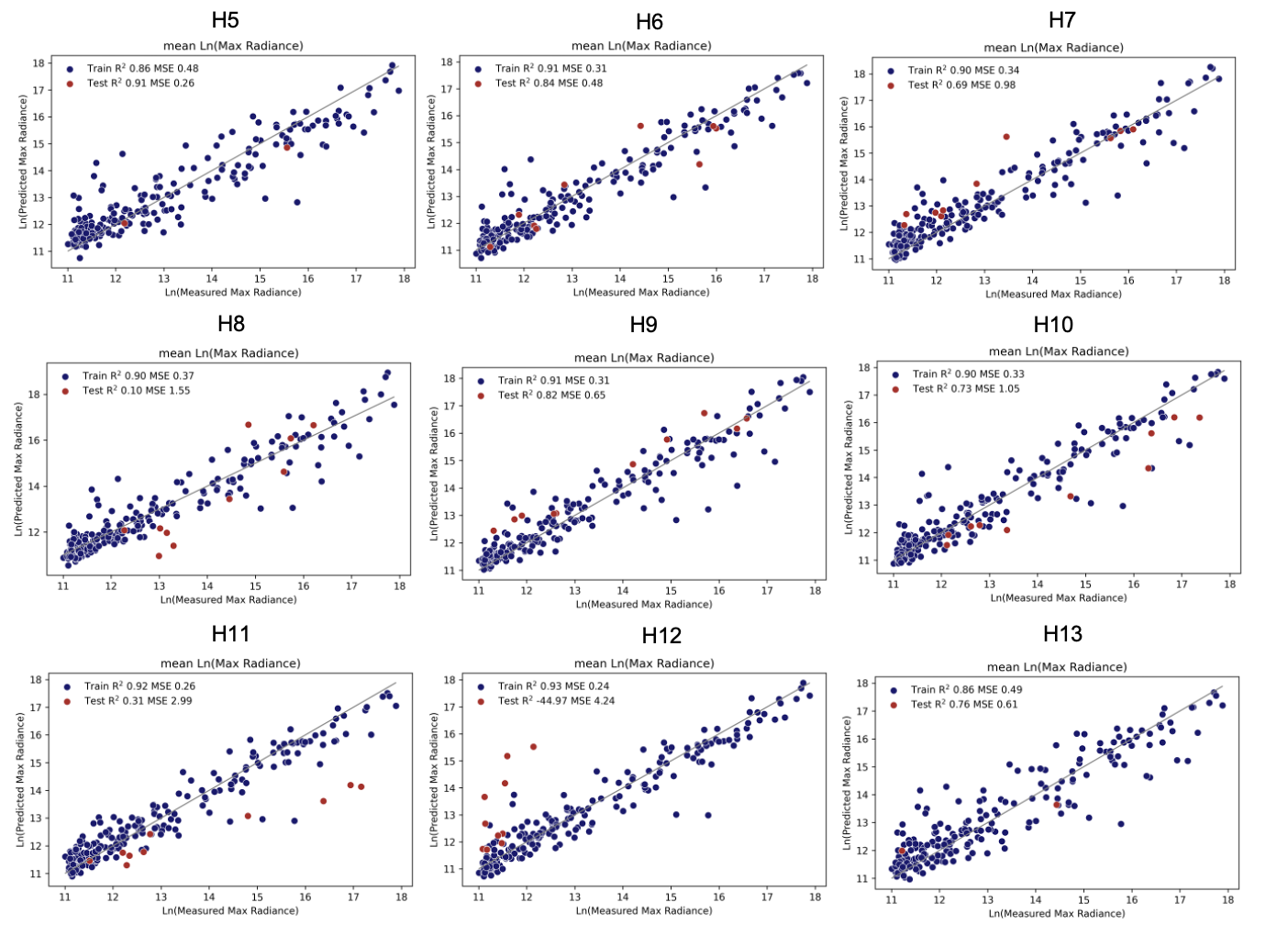

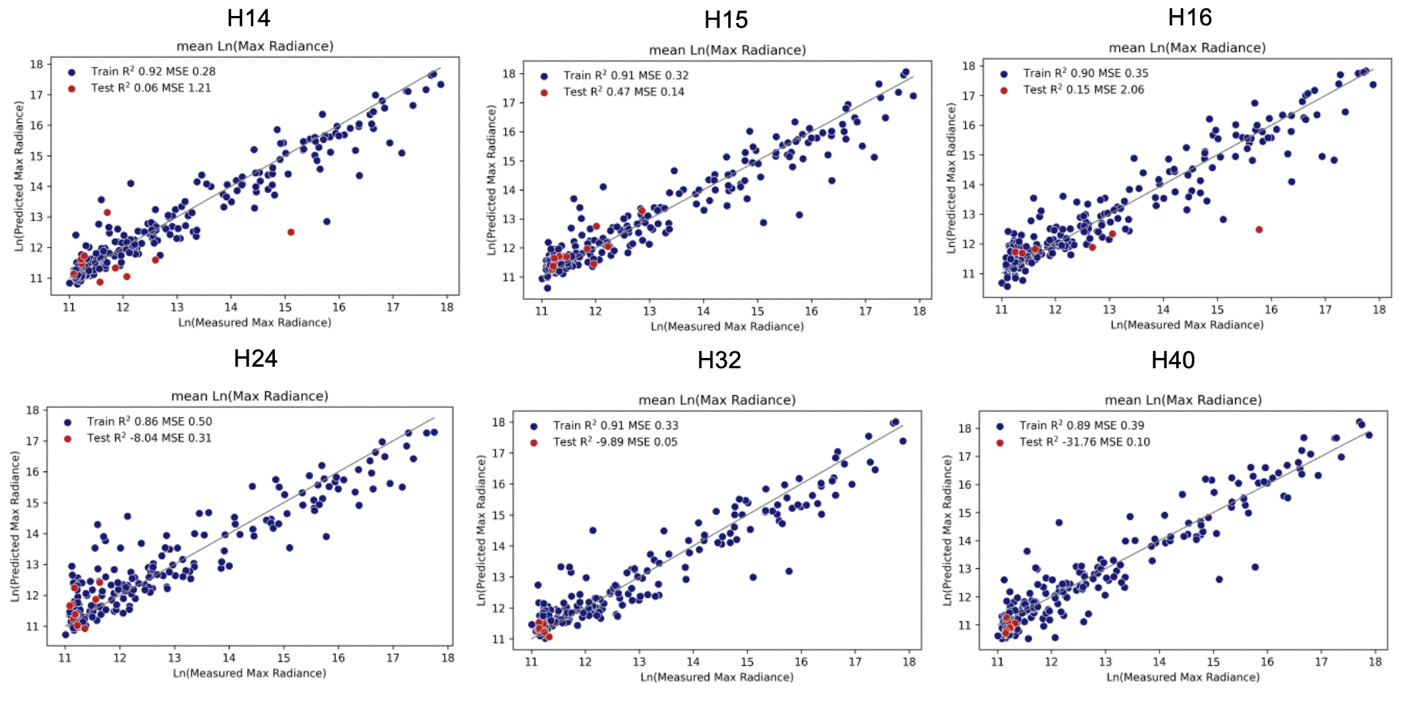

**Figure S4.** Leave-one-peptide-out cross-validation.

### Alternative Modelling Techniques

We compare our featurization method with two commonly used alternatives.

**1. Molecular Fingerprints**

Molecular fingerprints are a widely used, simple way to represent molecules by encoding the absence or presence of certain structural features. For each peptide, the full structure was first encoded into its respective SMILES string and converted to a Morgan circular fingerprint with 4096 bits and radius = 3.^5^ We reasoned that the large size of the extended peptides necessitated a large fingerprint size. Following fingerprint generation, all bits that did not vary for any of the peptides were removed, and the fingerprint concatenated with the peptide % and time to create the final feature array. Finally, random forest models were used to predict the minimum, mean, and maximum ln(max radiance) given the established superiority of tree-based methods for tabular data over neural networks.^6^ Here, hyperparameters were optimized using Bayesian optimization as implemented in scikit-optimize,^7^ however model performances did not exceed those obtained using default hyperparameters.

While moderate performance can be reached (Figure SXX), model performance is notably worse than our featurization method and GNN predictions.

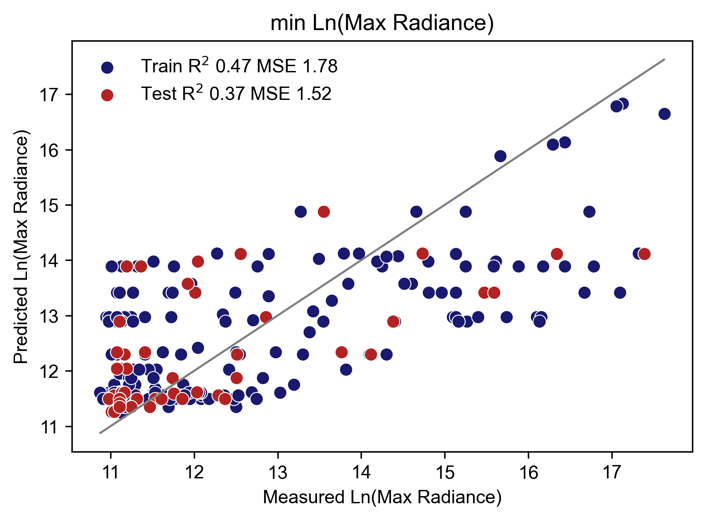

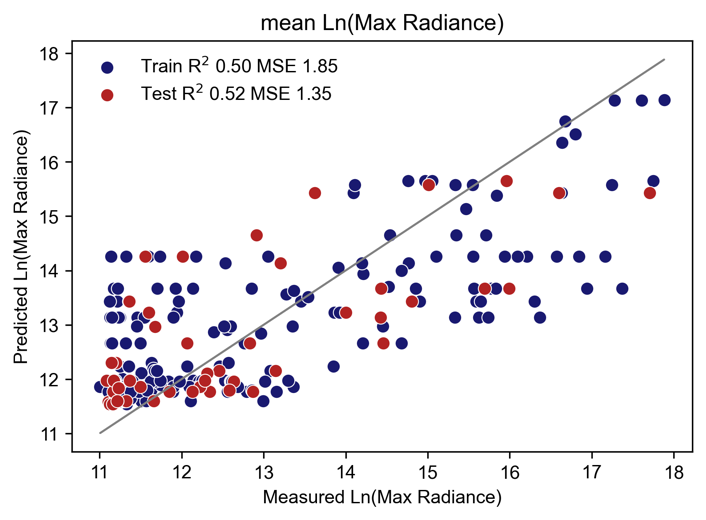

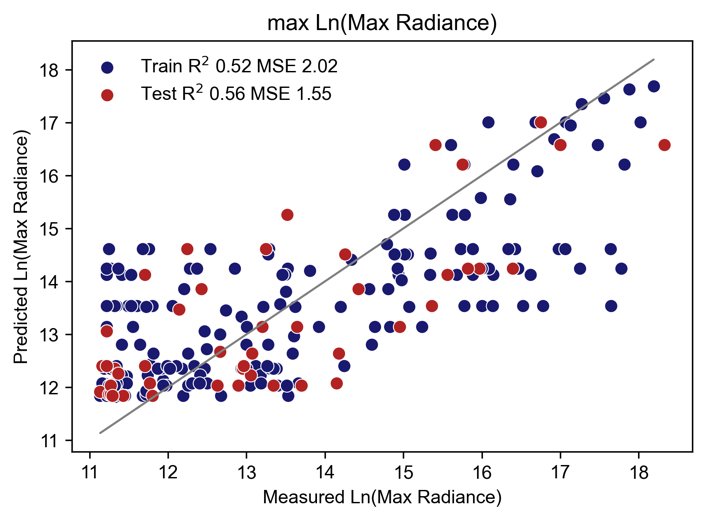

**Figure S5.** Benchmark models built with molecular fingerprints and random forest regressors.

2. Atomwise GNN

Alternatively, much success has been seen by treating atoms as nodes and bonds as edges. We use ChemProp,^8^ a widely used directed MPNN, to benchmark this approach. Like our approach, ChemProp uses a single layer MLP to provide a final prediction following aggregation operations. We refer the reader to supplementary reference^8^ for a full description of the ChemProp model. For settings shared with our approach, we deploy the same hyperparameters (i.e. epochs = 7000, no dropout, batch_size = 32, FFN hidden layer size = 514). Otherwise, we deploy ChemProp using its default settings. In all cases, poor model performance is observed.

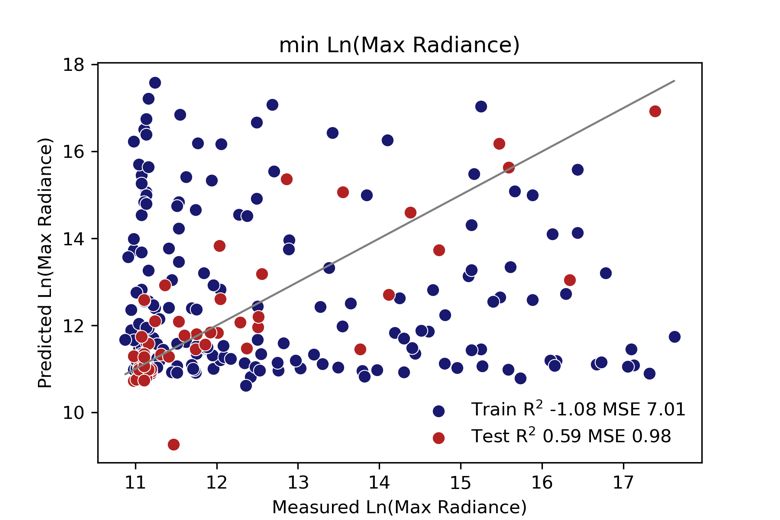

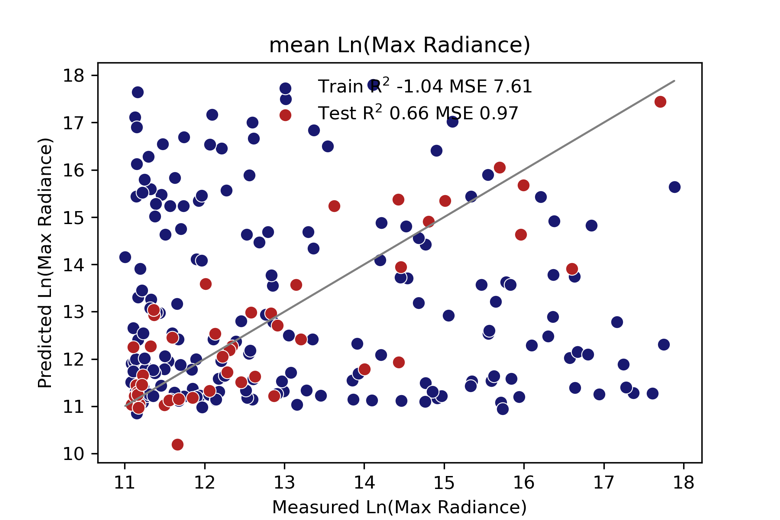

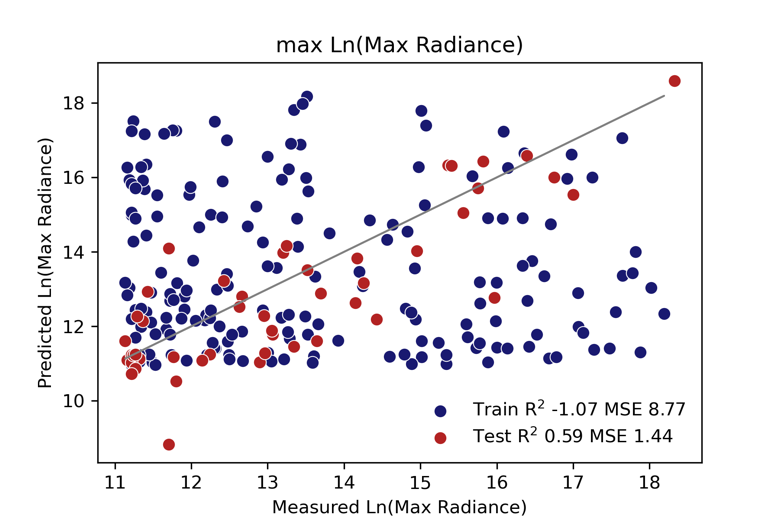

**Figure S6.** Atomwise GNN benchmarks.

### Virtual Screening

Prior to virtual screening, all available data was placed in the training set and the model retrained with 7000 epochs at the optimal hyperparameters. As mentioned above, variations in maximum radiance observed across different runs led to difficulties modelling all data at once. For virtual screening, we reasoned that separate models trained on the minimum, average, and maximum radiance would also provide greater insight than a single model. In other words, modelling these outputs separately should allow users to determine which peptides may be safe predictions (higher minimums) or those with higher ceilings (higher maximums). In practice, we find that the models trained on average values were much more noisy than those trained on maximum or minimum values. To rank prospective peptides in the main text, we use the model trained on maximum values. Models used for virtual screening are shown in Figure SXX.

**Virtual Screening Round 1**

**Figure S7.** Model used for virtual screening in round 1.

**Virtual Screening Round 2**

**Figure S8.** Models used for virtual screening in round 2.

### Peptide Legend

In the accompanying repository, peptides are generally denoted by the name SXX, or SXXR depending on the placement of protecting groups. Below we provide a visual legend for each peptide name (red = Ac, blue = H, yellow = K, grey = NH_2_) .

**Original Data**

**Virtual Screening Round 1**

**Virtual Screening Round 2**

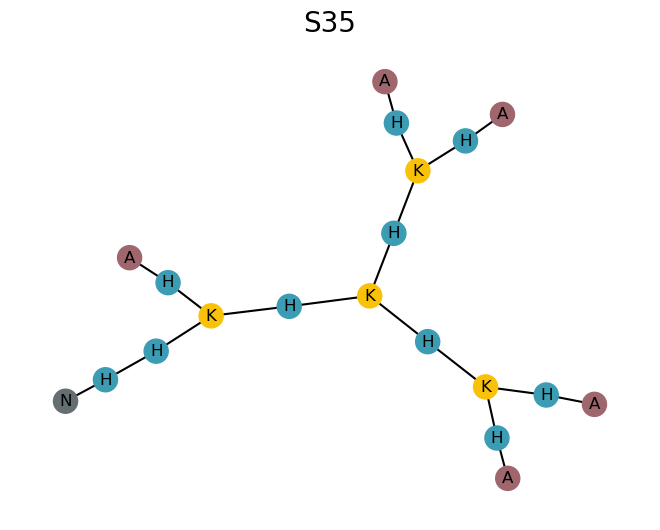

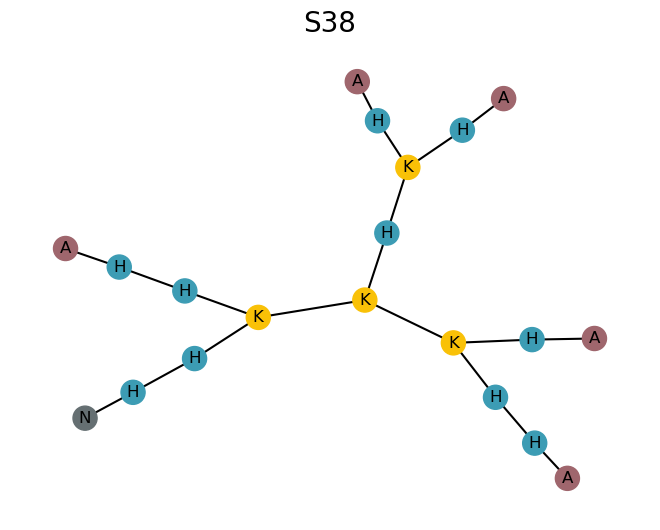

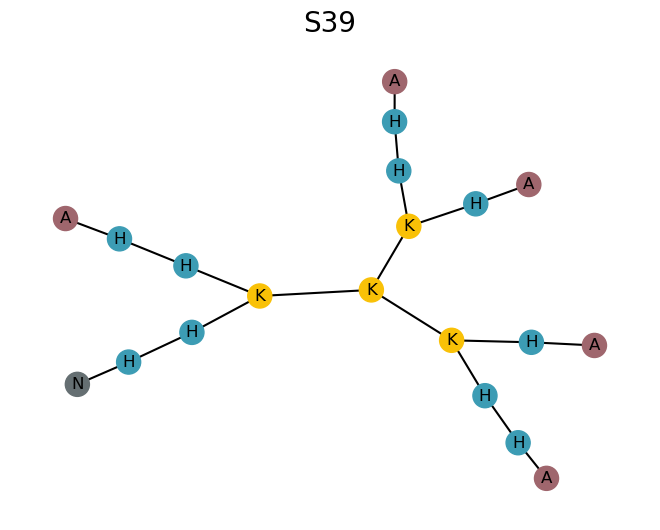
